# Prethalamus integrates neocortical and hypothalamic input to promote courage

**DOI:** 10.64898/2026.11.23.768550

**Authors:** Alice M. Koltchev, Sara Mederos, Alex Fratzl, Patricia Blakely, Nicole Vissers, Lucie Daveau, Sonja Hofer

## Abstract

Courage, the ability to act despite potential danger, is essential for adaptive behavior, yet the circuit mechanisms that balance caution and risk-taking under threat are not fully understood. Here we identify the ventral lateral geniculate nucleus (vLGN), an inhibitory prethalamic structure, as a key node in this process. High vLGN activity predicts courageous behavior, such as leaving a shelter to enter a potentially threatening environment, and optogenetic activation of vLGN neurons promotes risk-taking under diverse aversive conditions. Inputs to vLGN from retrosplenial cortex (RSC) and ventromedial hypothalamus (VMH) modulate this behavior in opposing directions: RSC input promotes leaving safety, while VMH input suppresses it. These inputs target largely distinct vLGN populations that interact to integrate these opposing drives: a hypothalamic signal promoting defensive caution and a neocortical signal promoting action despite potential threat.

## Introduction

Courage, the capacity to act despite potential danger, is essential for adaptive behavior. In uncertain or threatening environments, animals must balance avoiding harm with the need to explore and forage for resources. This requires a dynamic balance between caution and action: while excessive fear can prevent necessary action, suppressing fear entirely can lead to reckless or maladaptive behaviors.

Decades of research have identified circuits that govern fear and anxiety states, threat evaluation and defensive responses, involving widely distributed networks including amygdala, bed nucleus of the stria terminalis (BNST), hypothalamus, and brainstem areas such as the superior colliculus (SC) and the periaqueductal gray (PAG)^1–5^. On the other hand, circuits for appetitive motivation — including hypothalamic feeding and arousal centers, the mesolimbic dopamine system, cortico-striatal pathways and other circuits — drive reward-seeking, foraging and exploration^6–8^. In natural environments, however, animals rarely face purely defensive or appetitive challenges in isolation. Instead, they must arbitrate between competing motivational drives, weighing the potential benefits of a goal against the concurrent risk of harm^9–12^.

These decisions, historically often framed as approach-avoidance conflicts, require neural circuits that can integrate diverse contextual and sensory information with internal variables such as fear state, homeostatic needs and prior experience. Recent work has begun to reveal circuit mechanisms through which specific competing drives can influence approach-avoidance decisions. For instance, neurons and dopamine in the tail of the striatum regulate threat avoidance and learning to overcome it^13^, while distinct amygdalar, hippocampal and hypothalamic pathways bias behavior towards defensive responses or appetitive goals^14–20^. On top of this, neocortical areas can exert cognitive control, provide prior knowledge, promote exploration and enable adaptive coping^21–28^. Such affective and cognitive signals must ultimately be integrated to select one single behavioral outcome. Where this integration occurs remains unclear. Rather than residing solely within limbic or hypothalamic circuits that encode threat and internal state, it may involve structures that integrate these signals with cortical and sensory information to control behavioral output. The prethalamus is well positioned to perform such a role, receiving convergent cortical and subcortical inputs and influencing circuits that execute defensive behavior^29^.

In particular, the ventral lateral geniculate nucleus (vLGN), a predominantly inhibitory structure in the prethalamus, has been implicated in regulation of defensive behavior^29–31^. While its lateral part is densely innervated by the retina and can modulate the flow of visual information to downstream targets^32–38^, the vLGN’s role is not limited to visual processing. It receives rich and diverse inputs from many neocortical and subcortical areas and has been shown to be an important regulator of defensive reactions to threats and stress as well as influencing a wide range of other brain functions and behaviors, as diverse as motivation, feeding, memory and pain^30–32,39–45^.

In this study, we investigated if neural pathways through the vLGN could control mice’s decisions to leave a place of safety to explore a potentially threatening environment. We found that high activity of vLGN output to the midbrain predicts decisions to leave a safe space. vLGN activity is both necessary and sufficient to promote leaving safety across paradigms and can even reverse stress hormone–induced anxiety-like behavior. We then identified two excitatory inputs to the vLGN that contribute to these effects. Inputs from the retrosplenial cortex (RSC) and ventromedial hypothalamus (VMH) influence vLGN activity in opposite directions: activation of input from RSC promotes leaving safety, while input from VMH prevents it. These projections target distinct vLGN populations that interact locally, revealing a circuit mechanism by which competing cortical and hypothalamic drives are integrated within the vLGN to balance cautious and courageous behavior.

## Results

### Elevated vLGN activity promotes leaving safety

To investigate whether vLGN activity is associated with decision making under threat, we first recorded neural calcium dynamics using fiber photometry in mice expressing GCaMP6f in inhibitory vLGN neurons (**Figure 1A**). We imaged calcium activity of vLGN projections to intermediate and deep layers of the medial SC (mSC). The mSC is critically involved in defensive responses^46–49^, and projections from the vLGN to this structure have been shown to have strong control over defensive reactions to an imminent threat^30,31^.

**Figure 1.**
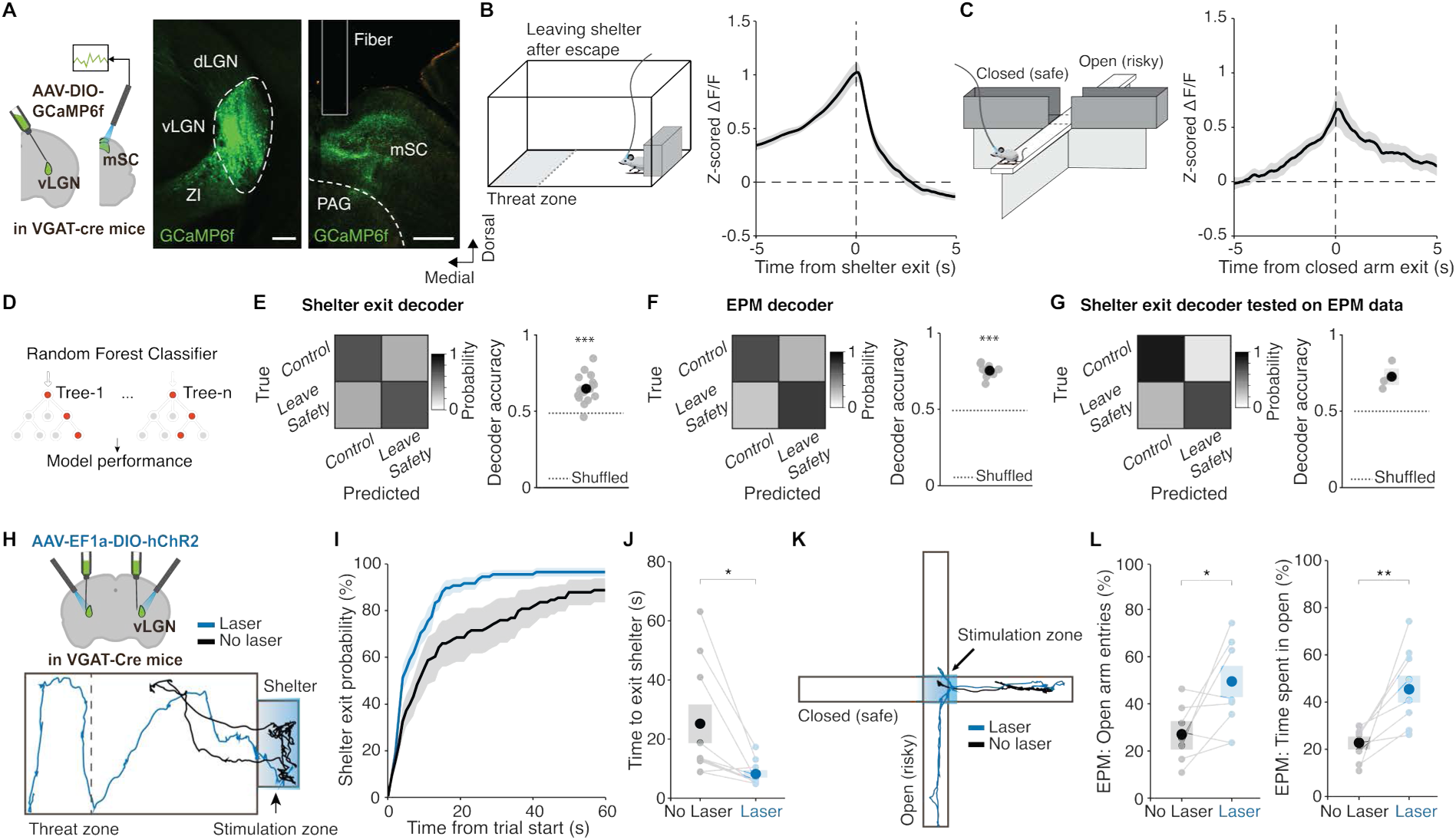
vLGN output activity predicts and causally promotes leaving safety across contexts. (A) Left, experimental approach. Middle, labelled GABAergic vLGN neurons, scale bar = 200 μm, and right, their axons in mSC, scale bar = 500 μm. dLGN: dorsolateral geniculate nucleus; mSC: medial superior colliculus; PAG: periaqueductal grey; ZI: zona incerta. (B) Left, schematic of the experimental arena with shelter. Overhead, looming stimuli are sporadically shown when mice enter the threat zone. Right, mean calcium activity of GABAergic vLGN axons in mSC aligned to animals leaving the shelter. N = 12 mice. (C) Left, schematic of the elevated plus maze (EPM). Right, mean calcium activity of GABAergic vLGN axons in mSC aligned to animals leaving a closed arm. N = 9 mice. (D) Decoding approach: A cross-validated random forest classifier was trained to predict shelter or closed arm exits from preceding vLGN calcium activity (-3s to 0s relative to exit). (E) Decoder performance trained and tested in shelter leaving paradigm. Left, mean row normalized confusion matrix for decoding shelter exits versus random control epochs (N = 12 mice). Right, decoder classification accuracy (p=0.0001), where each dot represents one animal, and the black dot and shaded box indicate mean ± SEM. (F) Decoder performance trained and tested in the elevated plus maze (EPM). Left, mean row normalized confusion matrix for decoding exits from closed arms versus random control epochs from vLGN calcium activity preceding closed arm exit (-3s to 0s) in the EPM (N = 9 mice). Right, decoder classification accuracy (p=0.0020; One-tailed Wilcoxon signed-rank test comparing observed decoding accuracy to a label-permutation null distribution n = 200 shuffles per animal, retrained classifier per shuffle). (G) Cross-paradigm generalization of decoding performance. Decoders trained on calcium activity in the leaving shelter paradigm were applied to vLGN activity of animals on the EPM, and performance was evaluated without retraining (N = 3 animals). Right, classification accuracy (p=0.1250). (H) Top, experimental approach. Bottom, representative mouse trajectories during 60s following trial start in the shelter during optogenetic activation of GABAergic vLGN neurons (blue) and in control (black) conditions. (I) Mean cumulative probability of animals’ exit from the shelter over 60s in trials with optogenetic activation of GABAergic vLGN neurons (blue) and control trials (black) within the same session. N = 10 mice. (J) Mean time until mice leave shelter after trial start during vLGN activation (blue) and in control trials (black). P = 0.015, two-tailed paired t-test, N = 10 mice. (K) Representative trajectories on the EPM during 30s following trial start after vLGN activation at the central choice point (blue) and in control trials (black). Light blue shading indicates the laser initiation zone. (L) Left, probability to enter open arms during laser stimulation at the central choice point (blue) and in control trials (black). P = 0.0241, two-tailed paired t-test, N = 8 mice. Right, percentage of trial time spent outside of closed arms after start of vLGN activation (blue) and in control trials (black). P = 0.0053, Welch’s t-test, N = 8 mice. In all plots, light dots represent data from individual mice. Error bars represent SEM across mice.

To determine whether vLGN also contributes to behavior beyond immediate threat responses, we examined its role in various paradigms. First, mice were placed in an elongated arena with a shelter at one end that the animals quickly adopted as a ‘safe place’^50^. When exploring the arena outside of the shelter, mice were occasionally presented with over-head threatening visual stimuli: expanding, dark (looming) disks that trigger escape back to the shelter^30,48,51^. After encountering such stimuli, mice remained in the safety of the shelter for some time, but eventually left this safe place on their own account, re-entering the potentially threatening environment. We found that calcium activity of vLGN axons in mSC strongly and consistently ramped up in the seconds preceding shelter departure (**Figures 1B, S1A**), suggesting that this circuit is engaged during internally driven decisions to re-enter a space associated with potential threat. This activity was modulated by animals’ threat expectation and could not be explained by their movements (**Figure S1C,D**) or the change in luminance^30^. Next, to assess whether these activity patterns generalize to other aversive contexts without imminent threat, we used the elevated plus maze (EPM), an anxiety-testing paradigm designed to leverage mice’s aversion to exposed places. The EPM features two “closed” arms surrounded by walls, and two “open” arms without walls that are innately aversive to mice (**Figure 1C**). Greater time spent in closed arms is used as a readout of heightened anxiety^52^, and the center of the maze represents a decision point at which animals need to leave the ‘safety’ of the closed arm if they want to explore the remainder of the maze. Consistent with the activity observed preceding shelter exits in the looming stimuli paradigm, vLGN axonal activity also ramped up prior to exits from the closed arms in the EPM (**Figures 1C, S1B**), suggesting a shared neural signature associated with transitions away from safety across contexts.

To formally test this, we trained random forest classifiers on peri-event activity (−3 s to 0 s) to decode transitions away from safety in each task. A cross-validated classifier trained in the leaving shelter paradigm could reliably identify leaving shelter events from preceding vLGN activity compared to random time periods within and even across animals (**Figures 1D,E, S2A,B**). Similarly, classifiers trained on EPM data could distinguish activity leading up to closed-arm exits from control periods (**Figures 1F, S2C,D**). Ramping activity leading up to leaving safety was thus distinctive and prominent enough to be highly predictive of exits from safety in both contexts. We then asked whether this representation generalized across tasks. Classifiers trained on vLGN activity preceding leaving shelter events in the looming arena could successfully identify when mice exited closed arms on the EPM (**Figure 1G**), indicating that a common neural representation in vLGN accompanies decisions to leave safety across distinct contexts and environments.

Given prior work showing that elevated vLGN activity suppresses defensive responses and promotes disengagement from threat^30,31,43^, these findings suggest that this signal may actively contribute to enabling mice to depart from safety into potentially dangerous environments. To test directly whether vLGN activity is causally involved in promoting such risk-taking behavior, we expressed channelrhodopsin (ChR2) in GABAergic vLGN neurons and delivered bilateral optogenetic stimulation in the same shelter-leaving assay (**Figure 1H**). We activated vLGN neurons in a subset of instances when mice were in the shelter for at least 5 seconds. This manipulation significantly increased the likelihood of mice exiting the shelter, making them leave the shelter approximately twice as fast as in control trials without laser stimulation (**Figures 1I,J, S3A,B**) and spend more time outside the shelter afterwards (**Figures S3C**). Laser stimulation without expression of optogenetic constructs had no effect on behavior (**Figure S3E-G**). In turn, when suppressing vLGN activity through systemic administration of CNO following bilateral expression of inhibitory DREADDs in GABAergic vLGN neurons, mice were less likely to leave the shelter and spent more time in it instead of exploring the arena (**Figures S3I-K**). This indicates that heightened vLGN activity is highly conducive to leaving safety.

To corroborate this effect across paradigms, we optogenetically activated inhibitory vLGN neurons in mice exploring the EPM. The laser was switched on only when animals reached the central choice-point from a closed arm, where they could decide to either cross over to the other ‘safe’ closed arm or explore one of the ‘risky’ open arms (**Figure 1K**). vLGN activation strongly increased the likelihood of mice to enter an open arm, and animals remained outside closed arms for longer afterwards (**Figure 1L**). These effects were strongest early in the session when mice were not yet habituated to the environment (**Figure S3D**) and not observed during laser stimulation in control mice not expressing ChR2 (**Figure S3H**). Chemogenetic suppression of vLGN activity in mice on the EPM made mice less likely to choose open arms at the decision point in the center^30^, particularly during the period when control mice first became more confident in exploring the open arms (**Figure S3L**). Together, these findings demonstrate that the level of vLGN activity plays a crucial role in determining the degree to which animals engage with or avoid aversive environments and demonstrate that heightened vLGN activity is sufficient to promote approach and exploration in the face of potential threat.

### vLGN activity promotes threat-discounting behavior

To further test the generality of vLGN’s role in courageous behavior, we manipulated vLGN activity in multiple other aversive contexts (**Figures 2, S4, S5**). We introduced a cricket-hunting paradigm to test the role of vLGN during more dynamic approach-avoidance conflicts. Mice can effectively hunt crickets, however, when first introduced, exhibit defensive behavior towards them, indicating that crickets are perceived as a potential threat^16,53^. When we activated vLGN during such a first encounter, mice approached the cricket more often, interacted with it for longer, and showed less defensive behavior (**Figures 2A-C; S4A-C**). Importantly, this additionally confirms that vLGN activation does not impair the animals’ vision, as prey pursuit, which is dependent on vision^54^, was not diminished but increased (**Figure S4B**). In a predator exposure task (**Figure 2D-F**), where mice could explore a room containing a toy snake, vLGN activation increased both the number of snake approaches and the time spent in its vicinity (**Figures 2D-F; S4D,E**). Importantly, this was only the case early in the session, before mice had habituated to the stimulus (**Figures 2E,F, S4E**). Together, these results indicate that vLGN activation reduces defensive avoidance and promotes approach despite potential danger. To test whether this is true only for visually-informed threats, we introduced an olfactory predatory threat (trimethylthiazoline, TMT, a component of fox feces) that induces strong defensive behavior in mice^55^, into the arena with the shelter (**Figure 2G**). Again, activating vLGN promoted faster departure from shelter (**Figure 2H**) and greater exploration of the arena (**Figure 2I**), confirming that the role played by vLGN in facilitating approach of potential threats is not specific to one sensory modality. Consistent with these results, and indicating that vLGN activity bestows positive valence, mice preferred the side that was paired with vLGN activation in a place-preference test (**Figures S4F,G**).

**Figure 2:**
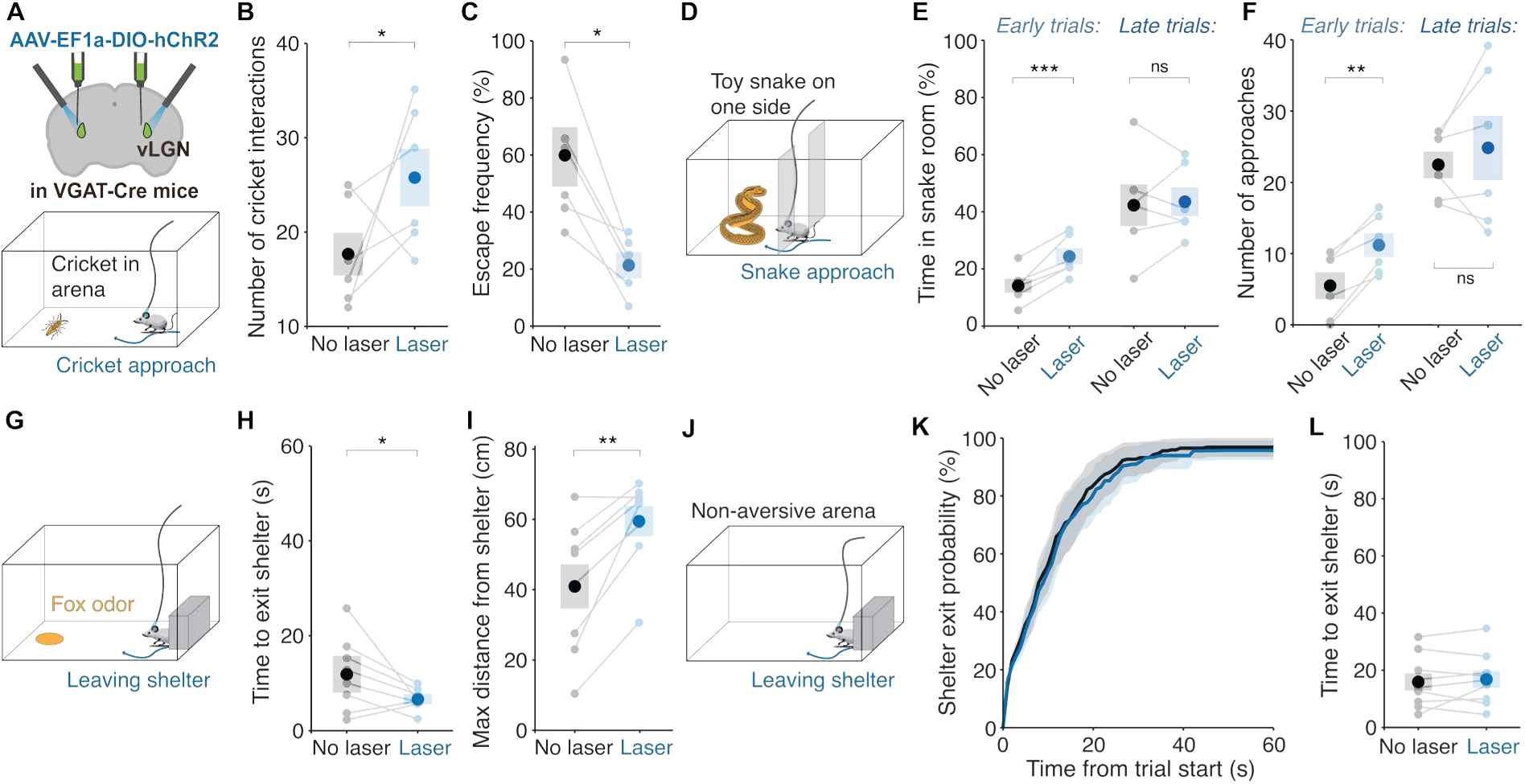
vLGN activation promotes threat-discounting behavior. (A) Cricket Test experimental protocol: ChR2-expressing GABAergic vLGN neurons are stimulated at 60s intervals for 30s in an arena with a living cricket. (B) Number of interactions with the cricket during vLGN activation (blue) and in control (black) periods. P = 0.0311, one-tailed paired t-test, N = 6 mice. (C) Escape frequency, as triggered by cricket approach or contact, during vLGN activation (blue) and during control periods (black). P = 0.0052, one-tailed paired t-test, N = 6 mice. (D) Snake Test experimental setup. ChR2-expressing GABAergic vLGN neurons are stimulated at 60s intervals for 30s. (E) Total time spent on the snake side early (first 10 minutes, left. P = 0.0005, one-tailed paired t-test, N= 6 mice) and late in the session (last 10 minutes, right. P = 0.4045, one-tailed paired t-test) when GABAergic vLGN neurons were optogenetically activated (blue) and in control trials (black) within the same session. (F) Same as H but for the number of snake approaches early (left, P = 0.0085, one-tailed paired t-test) and late (right, P = 0.2792, one-tailed paired t-test, N= 6 mice). (G) Experimental setup, with TMT (synthetic fox odour) in a petri dish away from the shelter. (H) Mean time to leave shelter after the start of activating ChR2-expressing GABAergic vLGN neurons in the shelter (blue) and in control trials (black). P = 0.0321, one-tailed paired t-test, N = 8 mice. (I) Maximum distance travelled away from the shelter during vLGN activation initiated in the shelter (blue) and in control trials (black). P = 0.0021, one-tailed paired t-test, N = 8 mice. (J) Schematic of arena with shelter without threat stimuli. (K) Mean cumulative probability of animals’ exit from shelter over 60s after start of activating ChR2-expressing GABAergic vLGN neurons in the shelter (blue) and in control trials (black). (L) Mean time to leave shelter after trial start. P = 0.3375, one-tailed paired t-test, N = 9 mice. In all plots, light dots represent data from individual mice. Error bars represent SEM across mice.

Next, we aimed to determine if effects of vLGN activation are specific to aversive context, or if it also affects exploration and general behavior in the absence of potential threat. First, we quantified shelter departures in a non-inversive environment with shelter. When animals had a long habituation time in the arena and had never experienced a threatening stimulus, vLGN activation had no effect on the likelihood or speed of animals leaving the shelter or their behavior once outside the shelter (**Figure 2J-L; S4H-J**). As further control experiments, we exposed mice to a sequence of items, either novel or familiar objects or food, one at a time. vLGN activation did not change the total time animals spent interacting with the objects when compared to control conditions (**Figures S5A-D**), nor the scale of their preference to novel compared to familiar objects (**Figure S5D**). In a two-novel-object choice paradigm, when vLGN was activated while mice interacted with one object while a second novel object was available, mice were not more likely to switch to the other object (**Figure S5E-G**), indicating that vLGN activation does not promote behavioral switching or state changes. Moreover, vLGN activation did not cause discernible changes in general behavior (**Figure S5H-J**). Together, these results show that while vLGN activation strongly suppresses defensive actions and promotes exploration in the face of various potential threats, its effects on behavior in all tested non-aversive contexts are negligible.

### vLGN activation counteracts stress-induced changes in defensive behavior

Decisions of whether to avoid or face potential threats are strongly influenced by internal states such as the overall anxiety level^10^. Thus, we next asked how modulation of internal state would affect mouse behavior in our paradigms and to what degree vLGN activation could counteract even strongly elevated levels of stress and anxiety. To this end, we gave mice a single intraperitoneal injection of the stress hormone corticosterone, whose receptors are widely expressed across the mouse brain and body, and which has been shown to induce a strong anxiety-resembling phenotype^56,57^. Thirty minutes after corticosterone injection, mice exhibited decreased exploration of exposed parts of an open field (**Figure 3A,B**) and spent significantly less time in the open arms of the EPM than saline-injected control mice (**Figure 3C**). Moreover, in the shelter-leaving paradigm, corticosterone-injected mice stayed in the shelter for longer after experiencing looming threat stimuli in the arena and spent less time outside it afterwards (**Figure S6A-C**). Remarkably, chemogenetically elevating vLGN activity via hM3D was sufficient to rescue this strong anxiety phenotype in all three paradigms (**Figures 3A-C, S6A-C**).

**Figure 3.**
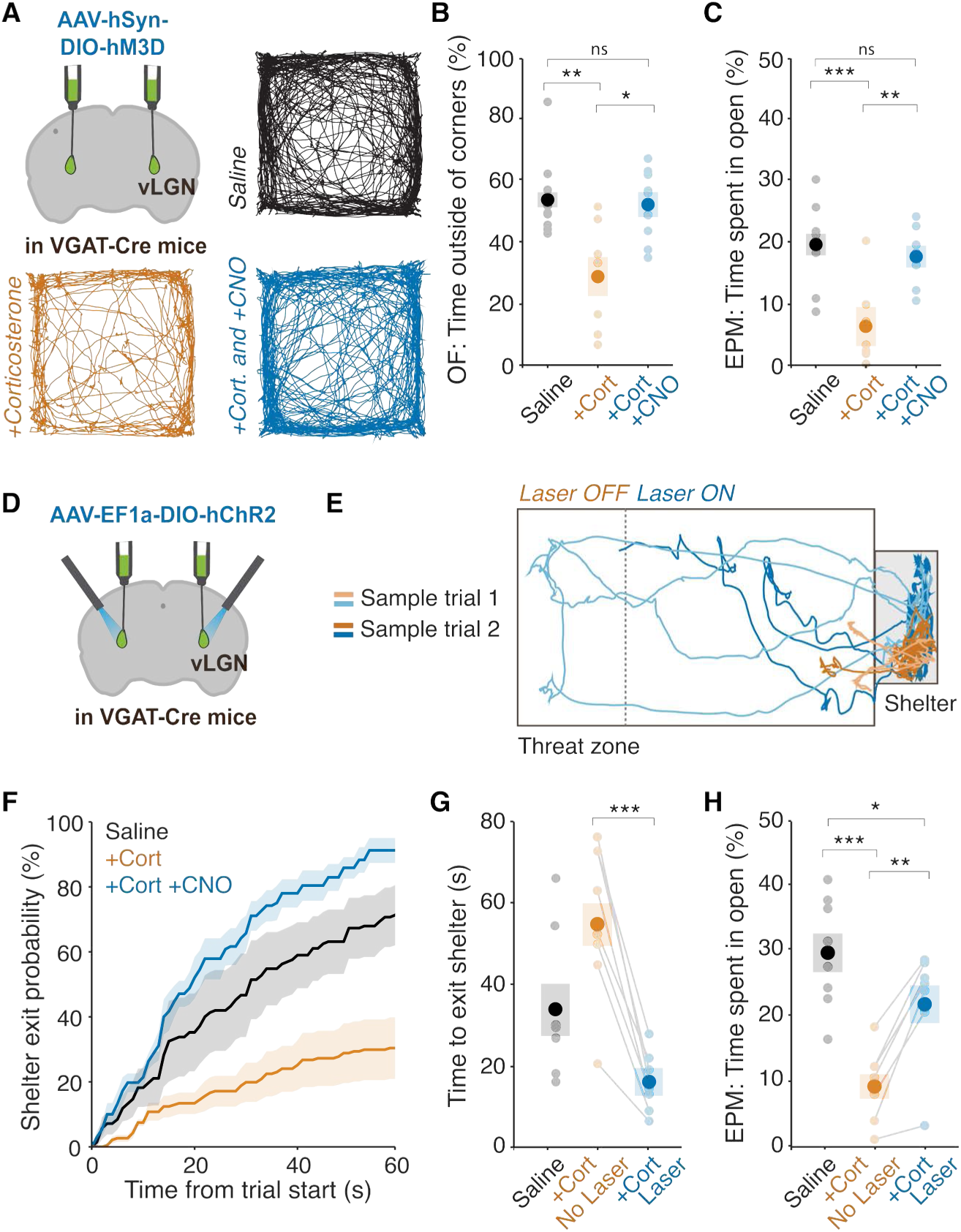
vLGN activation rescues stress-induced anxiety-like behavior. (A) Top left, experimental approach. Top right and bottom, representative trajectories over 60 s in an open field under different experimental conditions. (B) Percentage of time spent outside of the corners in an open field arena (OF) out of total session time for control mice injected with saline (black, N = 8 mice), mice injected with corticosterone (+Cort, orange, N = 8 mice), and mice injected with both corticosterone and CNO (+Cort+CNO, blue, N = 8 mice). One-way ANOVA, F(2,21) = 8.14, P = 0.002, followed by Tukey’s post hoc test: saline vs +Cort, P = 0.0028; saline vs +Cort+CNO, P = 0.72; +Cort vs +Cort+CNO, P = 0.017. (C) Percentage of time spent outside of the closed arms on an elevated plus maze (EPM) for saline (black, N = 8 mice), +Cort (orange, N = 8 mice), and +Cort+CNO (blue, N = 8 mice) mice. One-way ANOVA, F(2,21) = 13.49, P < 0.001, followed by Tukey’s post hoc test: saline vs +Cort, P = 0.0002; saline vs +Cort+CNO, P = 0.50; +Cort vs +Cort+CNO, P = 0.0027. (D) Experimental approach. (E) Representative trajectories of a +Cort mouse in two control trials (Laser OFF, orange) and two vLGN activation trials (Laser ON, blue) in the arena with shelter. (F) Mean cumulative probability of exit from shelter over 60 s from trial start for saline mice (black, N = 8 mice) and +Cort mice without (orange, N = 8 mice) and with (blue, N = 8 mice) vLGN activation in the same session. (G) Mean time to leave shelter after trial start for the same conditions as in F. Linear mixed-effects model with condition as fixed effect and animal as random intercept (Kenward-Roger degrees of freedom), F(2,11.6) = 43.9, P < 0.001, followed by Holm-corrected pairwise comparisons: saline vs +Cort No Laser, P = 0.058; saline vs +Cort Laser, P = 0.058; +Cort No Laser vs +Cort Laser, P = 0.0001. (H) Percentage of time spent outside the closed arms on an elevated plus maze for saline (N = 8 mice) and +Cort mice without and with vLGN activation (N = 8 mice). Statistics as in G: F(2,12.0) = 27.8, P < 0.001; saline vs +Cort No Laser, P < 0.0001; saline vs +Cort Laser, P = 0.010; +Cort No Laser vs +Cort Laser, P = 0.0012. In all plots, light dots represent data from individual mice. Error bars represent SEM across mice.

We then tested if brief vLGN activation could have a similarly penetrating effect. Optogenetic vLGN activation of corticosterone-injected mice in the shelter fully rescued behavior, increasing the animals’ likelihood to leave the shelter after threat experience above the level of control mice (**Figures 3D-G, S6D,E**). Similarly, optogenetic vLGN activation at the central choice-point of the EPM reversed behavior of corticosterone-injected mice, significantly increasing the time they spent in the open areas (**Figures 3H, S6F,G**). Thus, even brief vLGN activation robustly counteracted anxiety-like phenotypes across multiple contexts, despite corticosterone acting body-wide to induce a state of heightened stress. These findings establish the vLGN as a powerful regulator of defensive behavior: its activity is sufficient to override default defensive states and promote risk-taking, even under conditions of pathologically elevated anxiety.

### RSC input to vLGN drives exploration of aversive environments

Given the vLGN’s role in balancing cautious and courageous behavior, we set out to investigate which upstream brain areas contribute to this role through their projections to the vLGN. vLGN receives prominent layer 5 projections from visual^43,58^ (**Figure S7F**), but also higher-order cortical areas such as the anterior cingulate cortex (ACC) and retrosplenial cortex (RSC) (**Figures 4A, S7A**). We hypothesized that such corticofugal inputs could exert cognitive control over fear-related behavior through the vLGN. ACC in particular has been implicated in regulating fear, adaptive coping mechanisms, and effortful actions^23,27,59,60^. However, optogenetic activation of ACC projections in the vLGN had no measurable effect on the time it took mice to leave the shelter after experiencing a threat stimulus, or on their behavior on the EPM (**Figure S7A-E**).

**Figure 4.**
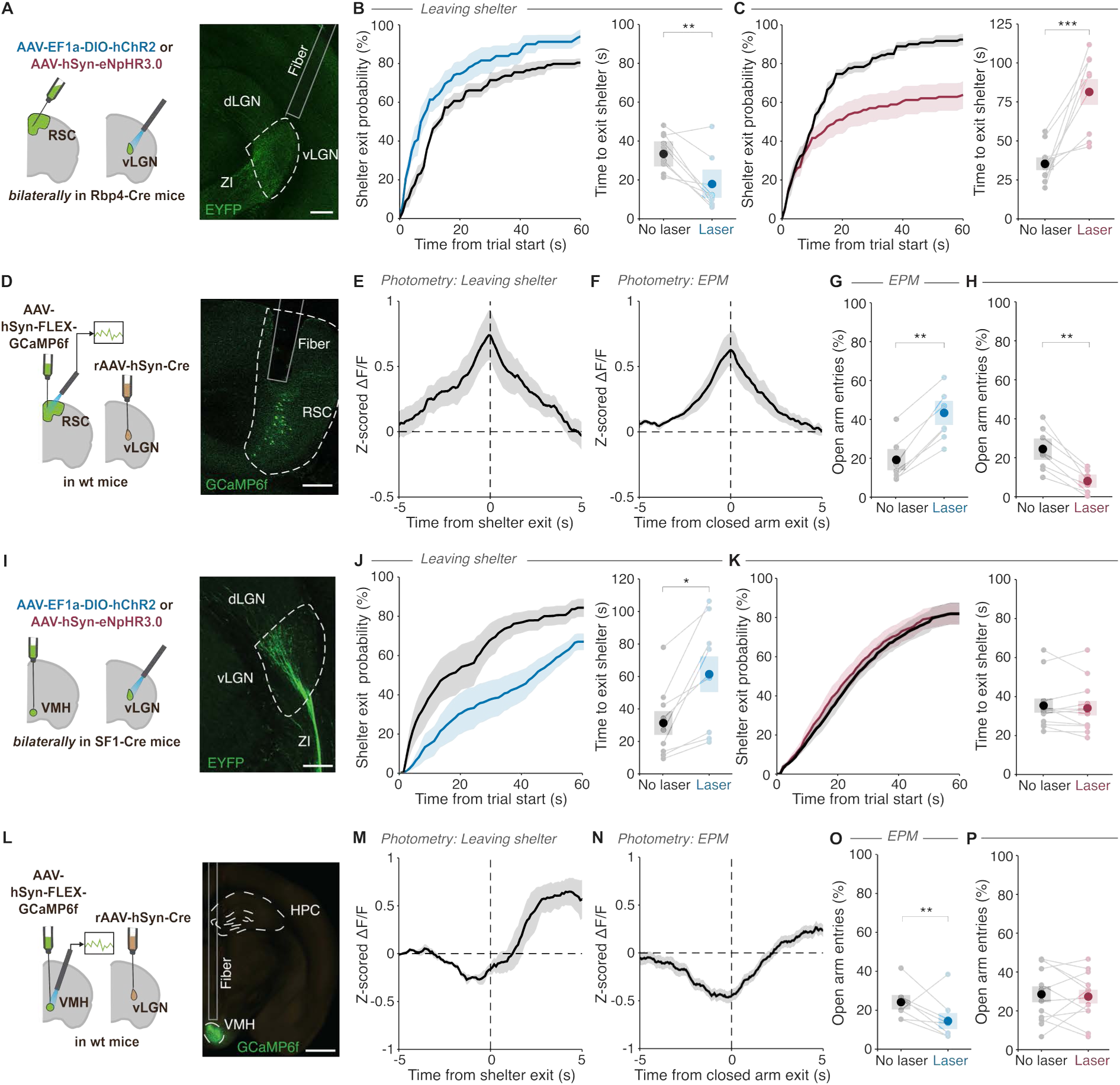
Retrosplenial cortex (RSC) and ventromedial hypothalamus (VMH) inputs to vLGN have opposing effects on leaving safety. (A) Left, experimental approach. Right, image of ChR2-labelled RSC axons in vLGN. Scale bar = 200 μm. (B) Left, mean cumulative probability of animals’ exit from shelter over 60s after start of optogenetic RSC axon activation in the shelter (ChR2 stimulation, blue) and in control trials (black). Right, mean time to leave shelter after trial start with (blue) and without (black) RSC axon activation. P = 0.0031, two-tailed paired t-test, N = 10 mice. (C) Left, mean cumulative probability of animals’ exit from shelter over 60s after start of RSC axon inactivation in the shelter (eNpHR3 stimulation, red) and in control trials (black). Right, mean time to leave shelter after trial start with (red) and without (black) RSC axon inactivation. P < 0.0001, two-tailed paired t-test, N = 9 mice. (D) Left, experimental approach. Right, image of vLGN-projecting RSC neurons. Scale bar = 200 μm. (E) Mean calcium activity from fiber photometry of vLGN-projecting RSC neurons aligned to shelter exit. N = 8 mice. (F) Mean calcium activity of vLGN-projecting RSC neurons aligned to animals leaving the closed arms of an EPM. N = 7 mice. (G) Percentage of time spent outside of closed arms after activation of RSC axons in vLGN at the central choice point of the EPM (ChR2 stimulation, blue) and in control trials (black). P = 0.0033, two-tailed paired t-test, N = 8 mice. (H) Percentage of time spent outside of closed arms after inactivation of RSC axons in vLGN (eNpHR3 stimulation, red) and in control trials (black). P = 0.0032, two-tailed paired t-test, N = 9 mice. (I) Left, experimental approach. Right, image of labelled VMH axons in vLGN. Scale bar = 200 μm. (J) Left, mean cumulative probability of animals’ exit from shelter over 60s after start of VMH axon activation (ChR2 stimulation, blue) and in control trials (black). Right, mean time to leave shelter, after the start of VMH axon activation (blue) and in control trials (black). P = 0.0382, two-tailed paired t-test, N = 9 mice. (K) Left, mean cumulative probability of animals’ exit from shelter over 60s after start of VMH axon inactivation (eNpHR3 stimulation, red) and in control trials (black). Right, mean time to leave shelter after start of VMH axon inactivation (red) and in control trials (black). P = 0.8349, two-tailed paired t-test, N = 12 mice. (L) Left, experimental approach. Right, image of vLGN-projecting VMH neurons. Scale bar = 500 μm. (M) Mean calcium activity from fiber photometry of vLGN-projecting VMH neurons aligned to shelter exit. N = 6 mice. Scale bar 1mm (N) Mean calcium activity of vLGN-projecting VMH somas aligned to animals leaving the closed arms of an EPM. N = 6 mice. (O) Percentage of open arm entries after VMH axon activation in vLGN at the central choice point of the EPM (ChR2 stimulation, blue) and in control trials (black). P = 0.0038, two-tailed paired t-test, N = 9 mice. (P) Percentage of open arm entries after VMH axon inactivation in vLGN at the central choice point of the EPM (eNpHR3 inactivation, red) and in control trials (black). P = 0.8297, two-tailed paired t-test, N = 9 mice. In all plots, dots represent data from individual mice and shading or error bars represent SEM across mice. dLGN: dorsolateral geniculate nucleus; HPC: hippocampus; RSC: retrosplenial cortex; VMH: ventromedial hypothalamus; ZI: zona incerta.

In contrast, the RSC→vLGN pathway had a clear influence on how animals acted in the face of potential threat. Optogenetic stimulation of ChR2-expressing RSC terminals in vLGN significantly decreased the time it took animals to leave the shelter after experience of threat stimuli in the arena (**Figure 4A,B**). Importantly, optogenetic suppression of the RSC→vLGN pathway specifically by laser stimulation of eNpHR3.0-expressing RSC terminals in vLGN impeded shelter exit (**Figure 4C**). Consistent with these findings, fiber photometry recordings of RSC neurons projecting to the vLGN revealed a prominent ramping signal prior to and peaking at shelter departure (**Figure 4D,E**), which could not be explained by animals’ motor actions (**Figure S8A,B**). The RSC→vLGN pathway therefore appears to be both necessary and sufficient for overcoming fear of potential threats. We corroborated these findings on the EPM, where RSC→vLGN calcium activity also ramped up before mice exited a closed arm (**Figure 4F**). Moreover, optogenetic activation of RSC inputs to vLGN at the central choice-point increased the likelihood of mice choosing the exposed open arms, while suppression of this pathway had the opposite effect and made mice more likely to choose the ‘safer’ closed arms (**Figure 4G,H**). We next tested if these effects are specific to RSC input, or if any input that activates vLGN circuits could promote threat-discounting behavior. Lateral visual cortical areas send dense projections to vLGN that can powerfully impact the activity of vLGN neurons and are necessary for learning to regulate fear reactions to visual threats^43^. However, optogenetic activation of lateral visual cortex input to vLGN had no effect on behavior on the EPM or in the leaving-shelter assay (**Figure S7F-J**). Together, these results highlight the RSC→vLGN projection as a functionally relevant neocortical input that supports courage-like decisions to overcome defensive states when threat is expected.

### The VMH-vLGN Pathway promotes risk-averse behavior

The vLGN also receives inputs from subcortical areas such as the hypothalamus (**Figure 4I**). The VMH, and in particular SF1-expressing neurons in this brain area, have been shown to encode a defensive state and predator imminence^61–63^, and high VMH SF1+ neuron activity promotes defensive behaviors such as freezing, escape, and avoidance in threatening contexts through multiple downstream targets^17,64,65^. We therefore investigated whether the VMH also modulates defensive behavior via a pathway through the vLGN. Anatomical tracing confirmed that VMH SF1+ neurons project to the vLGN (**Figures 4I**, see also **5A-D**). To test if and how this pathway influences behavior, we optogenetically activated VMH SF1+ axons in vLGN during the shelter-leaving paradigm. This manipulation had the opposite effect of RSC input activation; it suppressed shelter exit probability and significantly increased animals’ latency to leave the shelter compared to no-stimulation trials (**Figure 4J**). Similarly, activating this pathway in the center of the EPM strongly biased animals away from choosing the ‘riskier’, open arms (**Figure 4O**). These effects were specific to the VMH->vLGN pathway, because after ablation of vLGN neurons receiving VMH input, optogenetic activation of VMH axons in vLGN had no effect (**Figure S9**). To probe endogenous activity in this pathway, we recorded calcium signals from vLGN-projecting VMH somas (**Figures 4L-N**). Interestingly, VMH->vLGN activity decreased prior to shelter departure (**Figure 4M**) and prior to closed-arm exits in the EPM (**Figure 4N**), in contrast to the increase in activity before such risk-taking behavior apparent in RSC input to vLGN and vLGN output (**Figure 1B,C and 4E,F**). Together these data indicate that low VMH->vLGN drive is permissive for overcoming defensive states. However, optogenetic suppression of this pathway did not promote risk-taking behavior (**Figure 4K, P**), suggesting that lack of VMH input to vLGN alone is not sufficient to promote approach of potential threat, but that this necessitates a positive drive onto vLGN.

### Integration of competing drives in vLGN

The inputs from RSC and VMH into vLGN show opposing signals and have opposite effects on decisions around threat and safety. This suggests that the two pathways interact within vLGN to shape these decisions, either through convergent input or local circuit mechanisms. We found that RSC and VMH mostly target separate populations in vLGN with few vLGN neurons receiving input from both sources (**Figure 5A-D**). Importantly, calcium signals of vLGN neurons closely mirror the opposing activity patterns of their inputs—increased activity versus decreased activity before leaving safety in RSC-innervated versus VMH-innervated vLGN neurons, respectively (**Figure S8C-F**). Given that both RSC and VMH SF1+ projections are both glutamatergic and excitatory^66,67^, these different activity patterns in the vLGN further confirms that RSC and VMH drive distinct vLGN populations.

**Figure 5.**
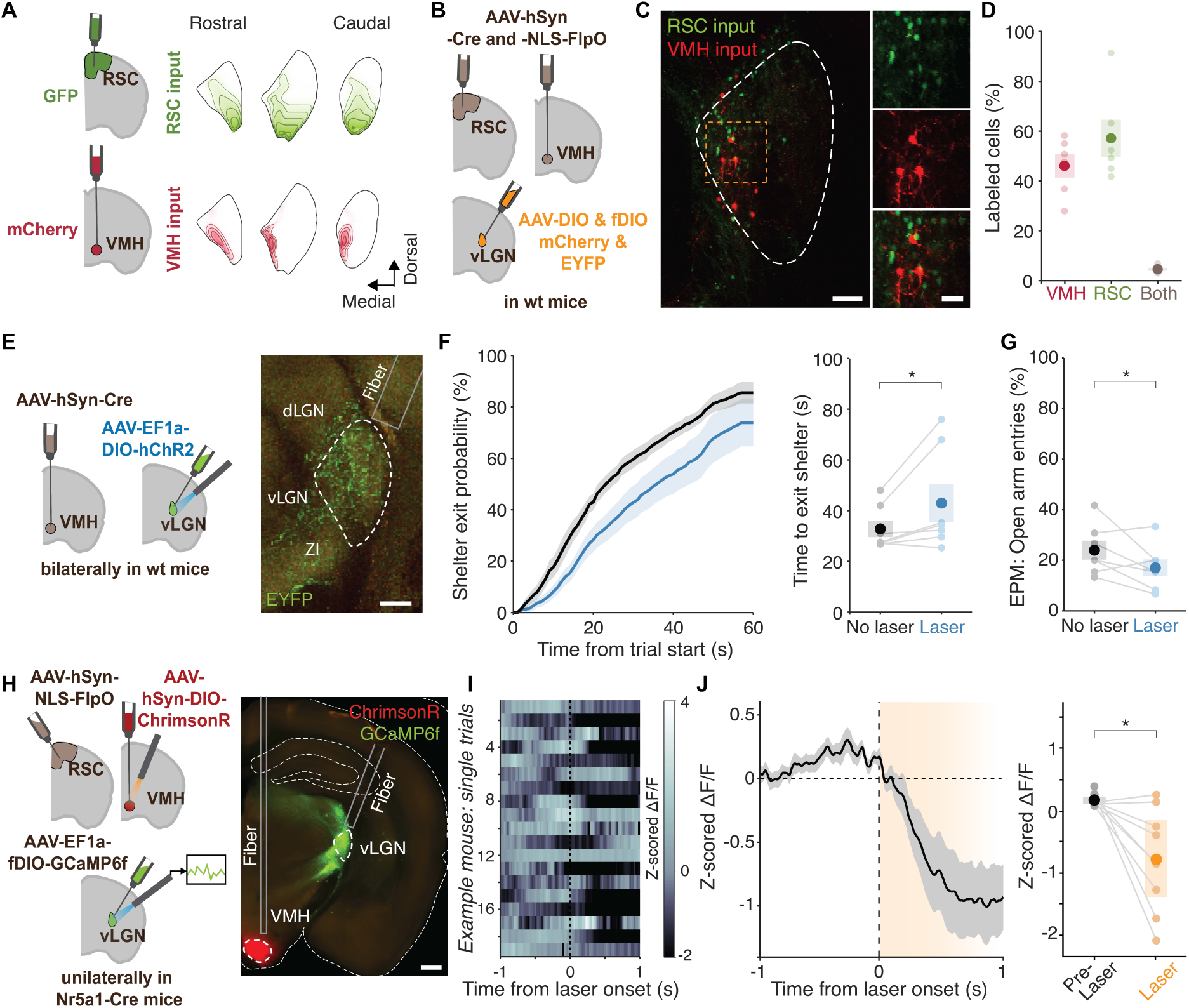
vLGN circuits integrate opposing cortical and hypothalamic drives. (A) Left, experimental strategy. Right, vLGN density maps averaged across five adjacent coronal slices and hemispheres, showing the distribution of labelled RSC and VMH axons across representative anterior, middle, and posterior levels of the vLGN, N = 3 mice. (B) Experimental strategy for labelling vLGN neurons innervated by RSC or VMH. (C) Representative confocal images showing labelled neurons in vLGN, Scale bar = 100 µm. Right, high magnification images. Scale bar = 10 µm. (D) Proportion of labelled vLGN cells innervated by VMH, RSC, or both. 5-9 slices per mouse, N = 6 mice. (E) Left, experimental strategy for optogenetic activation of VMH-innervated vLGN neurons. Right, representative image showing viral expression and optic fiber placement. Scale bar 200 µm. (F) Left, mean cumulative probability of animals’ exit from shelter over 60s after start of optogenetic activation of VMH-innervated vLGN neurons in the shelter (ChR2 stimulation, blue) and in control trials (black). Right, mean time to leave shelter after trial start with (blue) and without (black) activation of VMH-innervated vLGN neurons. P = 0.033, paired two-tailed t-test, N = 7 mice. (G) Percentage of open arm entries after activation of VMH-innervated vLGN neurons at the central choice point of the EPM (ChR2 stimulation, blue) and in control trials (black). P = 0.0435, paired two-tailed t-test, N = 7 mice (H) Left, Schematic of the combined optogenetic and fiber photometry strategy used to test whether VMH activation modulates activity in vLGN neurons receiving RSC input. Right, image of ChrimsonR-expressing VMH neurons and GCaMP6f-expressing RSC-innervated vLGN neurons. Scale bar, 500 µm. (I) Heatmap showing z-scored calcium activity from an example mouse across individual trials aligned to laser onset. (J) Average z-scored ΔF/F across mice aligned to laser onset. The orange shaded region indicates the laser stimulation period, the black line and grey shading show the mean and SEM. Right, statistics were computed by comparing mean activity before (-0.8 to 0 s) and after laser onset (0.2 to 1 s). P = 0.0116, paired two-tailed t-test, N = 9 mice. In all plots, dots represent data from individual mice and shading or error bars represent SEM across mice.

If high VMH activity promotes defensive behavior and cautious actions through a vLGN pathway, then activating VMH-innervated vLGN neurons should similarly promote defensive behavior, opposite to the effect of activating the general GABAergic vLGN population (**Figure 1I**). We found that this was indeed the case: optogenetic activation of vLGN neurons that receive VMH input slightly increased the time animals stayed in the shelter after threat experience and made them less likely to choose the exposed, open arms of the EPM (**Figure 5E-G**). One possibility of how this behavioral effect could be achieved is through local inhibitory interactions within vLGN, whereby VMH-innervated vLGN neurons inhibit those vLGN neurons whose activity promotes courageous decisions. To test this hypothesis, we combined activation of vLGN-projecting VMH neurons with imaging of RSC-innervated vLGN neurons (**Figure 5H**). Activity in these vLGN neurons was consistently suppressed by VMH stimulation, supporting inhibitory interactions between VMH and RSC input (**Figure 5I,J**). These interactions likely enable vLGN to either promote or prevent threat-discounting actions depending on the balance of those two inputs.

## Discussion

In this study, we identified the vLGN as a subcortical gate that integrates cortical and hypothalamic inputs to regulate decisions of whether to avoid or face potential threats. High neural activity of GABAergic vLGN neurons is crucial for promoting threat-discounting behavior across multiple aversive contexts – including in a model of pathologically heightened anxiety. Two excitatory inputs, from RSC and VMH, drive opposing behavioral effects through distinct vLGN populations that likely interact via local inhibition. These findings reveal a prethalamic circuit mechanism for integrating competing signals – likely containing information about the animal’s fear state from the hypothalamus^62,63^ and a neocortical drive to act in spite of it – to balance caution and courage.

The vLGN is a retinorecipient prethalamic structure of predominantly GABAergic neurons, classically assigned to the visual system^32^. However, its function goes beyond just visual processing. Several studies demonstrated an important role of the vLGN in the control of visually-driven defensive behaviors^30,31^, and different output pathways have been implicated in mediating the effect of bright light exposure on brain functions and behaviors as diverse as mood, vigilance, feeding, pain and itch^29,39,40,42,45^. These vision-related effects can mostly be attributed to retinorecipient neurons, often in the lateral (i.e. external) subdivision of the vLGN^34,38^. However, the vLGN also contains a medial (internal) subdivision that receives only sparse retinal input, but instead receives input from various cortical and subcortical sources, including the RSC and VMH. Our findings indicate that pathways through the vLGN can regulate fear and anxiety-related behavior across various sensory modalities, consistent with previous work that suggests a vision-independent role of vLGN circuits in stress resilience^44^.

There is no clear anatomical boundary between the vLGN and its lateral neighbor, the zona incerta. While minor spill-over of viral expression cannot always be prevented, we excluded animals with more significant viral expression in the zona incerta. Moreover, in a previous study, effects of neural vLGN manipulations were not reduced in animals with viral expression restricted to vLGN^30^. However, we cannot rule out that parts of the circuits relevant for courageous behavior described here reside in a border region of vLGN and very lateral zona incerta.

Our findings place the vLGN/ZI within a growing network of circuits that contribute to how animals resolve the competing demands of avoiding threat and engaging with the environment^13,14,16–19,24,27,28^. The involvement of several parallel pathways and distributed networks in these decisions is not surprising given their profound ethological importance. The ability to calibrate defensive behavior against other needs is among the most consequential computations an animal performs, and errors in either direction – excessive caution or reckless approach – carry severe survival risks. Different pathways may be preferentially recruited depending on the context, the nature of the threat, the animal’s motivational state, and the behavioral options available. The vLGN’s central location within these diverse pathways, downstream of neocortex and hypothalamus and upstream of brainstem and diencephalic structures involved in executing defensive actions, makes it well suited to serve as an integrative gate on defensive output. The architecture we propose, in which opposing drives converge onto anatomically distinct vLGN populations that interact through local inhibition, provides a mechanism for rapid, flexible integration of competing signals to regulate behavior.

One of these competing signals comes from the VMH, a central node in the hypothalamic predator defense system. SF1-expressing neurons in VMH in particular encode animals’ fear state and predator imminence, and their activation promotes freezing, escape, and avoidance through multiple downstream targets^17,61–65^. Here, we identify a parallel pathway through which VMH influences defensive behavior via the vLGN. VMH→vLGN activity decreased prior to departure from safety, and activation of this pathway suppressed threat-discounting behavior, while its silencing did not have a strong effect on its own. This asymmetry suggests that VMH input to vLGN functions mainly as a permissive brake: its withdrawal is important but not sufficient for courageous behavior, which additionally requires a positive drive – for instance in our experiments provided by the RSC. This positions the VMH→vLGN pathway as complementary to VMH projections to the PAG and the paraventricular thalamic nucleus, providing a mechanism through which the VMH can simultaneously promote defensive behavior via several pathways^17,65^, and suppress courageous actions via the vLGN.

We identified the RSC as a crucial input to vLGN to drive threat-discounting actions. The RSC is generally associated with spatial and contextual processing rather than fear regulation^68^. However, it is essential for spatial orientation during escape from threat^69,70^, plays a role in contextual fear memory storage and retrieval^71–73^ and is modulated by heightened stress^74^. Our finding that activity in RSC→vLGN projections is both necessary and sufficient to promote leaving safety under threat suggests that this pathway does not only transmit spatial information about shelter location^70^, but could also carry a signal that reflects the relative value of leaving the shelter despite potential danger. As such, the RSC may contribute a spatially informed "courage signal," combining knowledge of where safety is with a decision to leave it. Alternatively, RSC input could provide a more general drive to explore or gather spatial information about potential threats.

Despite previous work implicating ACC in fear regulation, cost-benefit decision-making under threat, and adaptive coping^21,23,27^, we found no effect of manipulating the ACC→vLGN pathway on behavior. However, this does not rule out a role of ACC in approach-avoidance decision-making. Recent work indicated that ACC contains distinct subpopulations with opposing functions – safety-encoding neurons that suppress defensive behavior via the zona incerta, and danger-encoding neurons that promote it via the SC^27^. Broad activation of ACC→vLGN axons may therefore simultaneously engage subpopulations with opposing functions that cancel each other out at the behavioral level. Or ACC→vLGN projections may be too sparse to drive behavior on their own, since the main projection of ACC to the prethalamus targets the zona incerta, not the vLGN^27^. ACC may thus exert an influence on threat-discounting behavior through other pathways. Alternatively, other medial prefrontal areas have been implicated in approach-avoidance decisions and may be more important for balancing caution and courage in the paradigms employed here^26,75^.

Our experiments were not designed to distinguish if vLGN activity suppresses an internal fear state or instead empowers animals to act despite fear. However, several points favor the latter interpretation. vLGN output pathways sit downstream of or parallel to regions important for the expression of fear and anxiety such as prefrontal cortical and hypothalamic areas, amygdala, and BNST^1–4,76^. The encoding of animals’ fear states is thus likely not strongly affected by vLGN manipulations. Moreover, even brief vLGN activation can fully and transiently counteract a systemic high-stress state induced by corticosterone injection. Receptors for this stress hormone are widely expressed^77^, causing whole-body effects that are unlikely to be fully reversed by brief vLGN activation. This suggests that vLGN circuits provide a behavioral bypass: rather than removing animals’ fear, it may allow them to act in spite of it and prioritize exploratory or appetitive drives over defensive behavior. This framing is consistent with theoretical models distinguishing fear regulation from fear tolerance, in which an organism retains awareness of threat but modulates the degree to which action is constrained^78–80^.

The balance between safety-seeking and engagement with a threatening world is fundamentally disrupted in anxiety disorders and PTSD, where patients exhibit excessive avoidance and an inability to act despite recognizing that avoidance is maladaptive^80,81^. Our finding that vLGN activation can override even pathologically heightened anxiety states suggests that prethalamic gating of defensive behavior may represent a locus at which this balance could be restored. While the primate homologue of vLGN, the pregeniculate nucleus, is much less well characterized, its conserved connectivity with brainstem and neocortex^32,82^ raises the possibility of targeting pregeniculate pathways to develop new approaches for the treatment of fear-related disorders.

## Acknowledgements

We thank Mehran Ahamadlou and Thomas Mrsic-Flogel for feedback on the manuscript and Mrsic-Flogel and Hofer Lab members for helpful discussions. We thank M. Li for animal husbandry and genotyping; R. Campbell, J. Broni-Tabi, and the Sainsbury Wellcome Centre (SWC) imaging core for help with serial section two-photon imaging and slide scanner imaging; P. Nowak from the SWC viral core for providing viruses. We thank the SWC FabLabs for technical support. This work was supported by the Sainsbury Wellcome Centre core grant from the Gatsby Charitable Foundation and the Wellcome Foundation (090843/F/09/Z), a Wellcome Investigator Award (S.B.H., 219561/Z/19/Z), an EMBO postdoctoral fellowship (S.M., EMBO ALTF 327-2021), a Wellcome Early Career Award (S.M., 225708/Z/22/Z), a La Caixa Junior Leader fellowship (S.M., 160957), and a Ramón y Cajal Program award (S.M., RYC2024-049051-I).

## Author contributions

A.F., A.K., S.M. and S.B.H. conceived the study and designed the experiments. A.F. and S.M built the experimental setups. A.K., S.M and A.F. performed the experiments, and analyzed the data, with help from P.B., N.V., and L.D.. A.K., S.M. and S.B.H. wrote the paper, with inputs from A.F. and N.V..

## STAR Methods

### Experimental model and subject details

Mice were housed in individually ventilated cages (IVC) under controlled climate (temperature 20-24°C, humidity 45-65%) in a reversed light:dark cycle (12:12h) with free access to food and water. 8-16 week old male and female VGAT-ires-Cre mice (Vong et al., 2011, Jackson Laboratory, stock 028862), Nr5a1-Cre mice, also called SF1-Cre mice (Dhillon et al., 2006, Jackson Laboratory, stock 037533), Rbp4-Cre mice (Gong et al., 2007, MMRC, stock 031125-UCD), and wild-type (WT) C57BL/6J mice (Charles River) were used for the experiments. Data from male and female mice were pooled. All experiments were performed in accordance with UK Home Office regulations (Animal Welfare Act of 2006) under project license (PPL) PD867676F, following local ethical approval.

### Viruses

Viruses used in this study are listed here and referred to by abbreviations in the main text. Viruses 6 - 12 were produced in-house by the SWC virus core.

1. AAV1-EF1α-double-floxed-hChR2(H134R)-EYFP-WPRE-HGHpA (ChR2; Addgene 20298-AAV1; provided by Karl Deisseroth) was used for optogenetic stimulation at 7.0 × 10¹² vg ml⁻¹.
2. AAV1-hSyn-DIO-ChrimsonR-tdTomato (provided by the SWC Viral Vector Core Facility) was used for optogenetic stimulation at 3.7 × 10¹² vg ml⁻¹.
3. AAV2-EF1α-DIO-eNpHR3.0-mCherry (UNC Vector Core) was used for optogenetic silencing of axons in the vLGN at final titres ranging from 4.0 × 10¹¹ to 2.0 × 10¹² vg ml⁻¹.
4. AAV1-Syn-Flex-GCaMP6f-WPRE-SV40 (GCaMP6f; Addgene 100833-AAV1; provided by Douglas Kim and the GENIE Project; Chen et al., 2013) was used for calcium imaging at 2.81 × 10¹² vg ml⁻¹.
5. AAV1-hSyn-DIO-hM4D(Gi)-mCherry (hM4Di; Addgene 44362-AAV1; provided by Bryan Roth; Krashes et al., 2011) was used for chemogenetic inhibition at 9.0 × 10¹² vg ml⁻¹.
6. AAV1-hSyn-DIO-hM3D(Gq)-mCherry (hM3Dq; Addgene 44361-AAV1; provided by Bryan Roth; Krashes tet al., 2011) was used for chemogenetic excitation at 9.0 × 10¹² vg ml⁻¹.
7. AAV1-EF1α-FLEX-EGFP was used for chemogenetic control experiments and anatomical tracing at 2.0 × 10¹³ vg ml⁻¹.
8. rAAV2-retro-hSyn-Cre was used to express Cre recombinase in neurons projecting to the vLGN at 1.0 × 10¹⁴ vg ml⁻¹.
9. rAAV2-retro-EF1α-FlpO Cre was used to express Flp recombinase in neurons projecting to the vLGN at 3.55 × 10¹⁴ vg ml⁻¹.
10. AAV1-hSyn-NLS-FlpO was used for anatomical cell quantification and calcium imaging at 3.43 × 10¹⁴ vg ml⁻¹.
11. AAV1-EF1α-fDIO-GCaMP6f was used for calcium imaging at 7.10 × 10¹⁴ vg ml⁻¹ and for anatomical tracing at 3.55 × 10¹⁴ vg ml⁻¹.
12. AAV1-EF1α-FLEX-tdTomato was used for anatomical cell quantification at 5.0 × 10¹³ vg ml⁻¹.
13. AAV5-EF1α-Flex-taCasp3-TEVp was used to ablate vLGN neurons receiving input from the VMH at 3.6 × 10¹⁴ vg ml⁻¹.
14. AAV1-CaMKIIα-hChR2(H134R)-EYFP was used for optogenetic activation of VMH axons in the vLGN at 2.3 × 10¹⁴ vg ml⁻¹.

### Surgical procedures

Prior to surgery, the analgesic Metacam (5 mg/kg) was administered subcutaneously. General anaesthesia was induced (4%) and maintained (1-2.5%) with Isoflurane (in oxygen, 1L/min flow rate). Body temperature was maintained at 36.5°C using a controlled heating pad. The eyes were protected from light and drying by application of Xailin lubricating eye ointment. Mice were head-fixed on a stereotaxic device (Kopf, model 940) using ear bars. The skull was exposed by using a scalpel to cut along the midline. Craniotomies were made using a dental drill (Foredom rotary micromotor) and viral vectors were delivered using pulled glass pipettes (4.2uL glass capillaries with ID 0.53nm, Drummond Scientific) in an injection system coupled to a programmable nanoliter injector (Nanoject III, Drummond Scientific) at approximately 4 nL/min. Pipettes were sharpened using a microgrinder (EG-45, Narishige). Ten minutes after the injection, the pipette was retracted and the skin was either glued (Vetbond), or optic fibers were implanted. Optic fibers were inserted 100 μm (for optogenetics) or 50 μm (for photometry) above the target area and affixed using light-cured dental cement (RelyX Unicem 2, 3M). Animals were provided with wet calorie-dense food and Metacam jelly for 48 h post-surgery. Optogenetic manipulation and photometry experiments were performed 14 to 35 days post-surgery. Chemogenetic manipulation experiments were performed 28 to 35 days post-surgery.

For most experiments, viruses were injected bilaterally in the vLGN (40-65 nL per side, from Bregma: ML: ± 2.52 mm, AP: −2.3 mm, DV: −3.45 mm). For optogenetic manipulation experiments, two 200 μm optic fibers (MFC_200/245-0.37_4.5mm_ZF1.25_FLT, Doric Lenses) were implanted bilaterally over vLGN without damaging the optic tract (from skull at Bregma: ML: ±2.6 mm, AP: -2.1 mm, DV: -3.18 mm, 20 degrees lateral to medial). For fiber-photometry recordings of vLGN cell bodies, a single 400 μm optic fiber (MFC_400/475-0.53_4.5mm_ZF1.25_FLT, Doric Lenses) was implanted over vLGN without penetrating the optic tract (from skull at Bregma, ML: ±3.5 mm, AP: -2.31mm, DV: -2.12 mm, 20 degrees lateral to medial).

For RSC fiber-photometry, optogenetic, and anatomical experiments, viruses were injected unilaterally or bilaterally in the RSC of WT (combined with retroAAV injection in vLGN) or Rbp4-Cre mice (120 nL per injection, at two sites, from skull at Bregma: ML: ± 0.5 mm, AP: -2.7 mm, DV: between -0.9 and -1.5 mm, and ML: ± 0.7 mm, AP: -3.2 mm, DV: between -0.8 and -1.4 mm). For optogenetic manipulation experiments, two 200 μm optic fibers (MFC_200/245-1.66_2mm_ZF1.25_FLT, Doric Lenses) were implanted bilaterally over RSC (from skull at Bregma: ML: ± 0.6 mm, AP: -2.95 mm, DV: between -1 mm). For fiber-photometry recordings of RSC cell bodies, one 400 μm optic fiber (MFC_400/430-1.66_2mm_ZF1.25_FLT, Doric Lenses) was implanted over RSC (from skull at Bregma: ML: +0.6 mm, AP: -2.95 mm, DV: -1 mm).

For ACC fiber-photometry, optogenetic, and anatomical experiments, viruses were injected unilaterally or bilaterally in the ACC of WT mice (combined with retroAAV injection in vLGN) for fiber photometry experiments or Rbp4-Cre mice for optogenetic manipulation of axons in vLGN (120 nL per injection, at two sites, from skull at Bregma: ML: ±0.35 mm, AP: 1 mm, DV: between -1.9 and 2.1 mm, and ML: ±0.35 mm, AP: 0.5 mm, DV: between -1.6 and -1.9 mm). For optogenetic manipulation, two 200 μm optic fibers (MFC_200/245-1.66_2mm_ZF1.25_FLT, Doric Lenses) were implanted bilaterally over vLGN.

For VMH photometry and optogenetic experiments, viruses were injected unilaterally or bilaterally in the VMH of wild-type or Sf1-Cre mice for optogenetic manipulation of axons in vLGN (70 nL per injection, from skull at Bregma: ML: ± 0.5 mm, AP: -1.31 mm, DV: -5.6 mm). For optogenetics experiments, two 200 μm optic fibers (For fiber-photometry recordings of VMH cell bodies, one 400 μm optic fiber (MFC_400/430-1.66_6mm_ZF1.25_FLT, Doric Lenses) was implanted over VMH (from skull at Bregma: ML: ± 0.5 mm, AP: -2.31 mm, DV:-5.45 mm). For photometry, or two 200 μm optic fibers (MFC_200/245-0.37_6mm_ZF1.25_FLT, Doric Lenses) were implanted bilaterally over VMH (from skull at Bregma: ML: ± 0.5 mm, AP: -2.31 mm, DV: -5.45 mm) or over vLGN.

### Histology

At the end of experiments mice were euthanized with an intraperitoneal dolethal injection (IP, 80 mg/kg) and transcardially perfused (0.01 M phosphate buffered saline (PBS), followed by 4% paraformaldehyde in PBS). For anatomical confirmation of injection sites, fiber placement and visualization of labelled neurons and axonal projections, brains were embedded in 5% agarose (A9539, Sigma) and imaged using a custom-built serial-section two-photon microscope (Mayerich et al., 2008; Ragan et al., 2012). Coronal slices were cut at a thickness of 50 μm using a vibratome (Leica VT1000), and optical sections were acquired every 12.5 μm. Scanning and image acquisition were controlled by ScanImage v5.5 (Vidrio Technologies, USA, Pologruto et al., 2003) using a custom software wrapper for defining the imaging parameters (https://zenodo.org/record/3631609).

To quantify the proportion of RSC-, VMH-, and ACC-innervated neurons in vLGN, coronal slices were kept in 0.01M PBS after imaging for up to 24 hours before being mounted with DPX mountant (06522, Sigma Aldrich). Images were acquired using a slide-scanner (Axioscan, Zeiss). The proportion of RSC-, VMH-, and ACC-innervated neurons in vLGN was quantified manually on single-plane images using Fiji, a wrapper for ImageJ^83^.

### Experimental setups

Experiments assessing leaving shelter after exposure to looming stimuli were performed in a custom-made transparent acrylic arena (“escape arena” L: 80 cm, W: 26 cm, H: 40 cm; Fratzl et al., 2021). The opaque floor was white to improve automated tracking. A custom-made, red-tinted acrylic shelter (L: 14 cm, W: 14 cm, H: 14 cm) was placed on one end of the arena. A small empty Petri dish was anchored at the other end of the arena to promote exploration. Experiments assessing leaving shelter in the presence of an olfactory predatory stimulus, TMT (2,4,5-Trimethylthiazole, 219185, 1.013 g/mL, Sigma-Aldrich) was placed on a cotton bud inside the Petri dish. Cricket interactions were assessed in an arena of the same dimensions, empty apart from a live cricket (Black Female Adult crickets, Blades Biological). Experiments assessing approach behavior in the context of a toy snake (Toyandona) and place preference experiments were performed in an identical acrylic arena with two red acrylic panels dissecting the long axis of the arena, forming two separate equally-sized chambers with a narrow opening.

The open field test was performed in a custom-made white acrylic arena (L: 40 cm, W: 40 cm, H: 30 cm). All novel-object tests were also performed in the same acrylic arena. The elevated plus maze (EPM) consisted of two opposing open arms (without walls, L: 30 cm, W: 5 cm) and two opposing closed arms (with walls, L: 30 cm, W: 5 cm, wall H: 5 cm). All four arms extended from a center square (5 cm by 5 cm) to form a plus shape. The maze was positioned on an acrylic pedestal (H: 20 cm).

All arenas were placed in a large light-proofed and sound-deadening box (L: 120 cm W: 100 cm H: 120 cm, LS Fabrications). An ultra-short throw DLP projector (HF85JA, LG) was mounted on an custom-made aluminum scaffold (Bosch Rexforth) within the box and back-projected via a mirror onto a suspended horizontal screen (60 cm above arena floor, 100 cm 3 80 cm; ‘100 micron drafting film’, Elmstock) For all behavioral assays, the screen was kept at a constant background luminance level of 9 cd x m^-2^. Arenas were also illuminated by a strip of infrared LED illuminators (Jcheng Security) to ease tracking of the mice. Behavior was recorded with a near-IR GigE camera located above the arena (acA1300-60 gmNIR, Basler) at 50 frames per second through a varifocal lens (H2Z0414C-MP, Computar) and 780 nm long-pass filter (Thorlabs).

In one setup (setup A), camera frames (50fps) were triggered and synchronized by the LabVIEW-based (National Instruments) Mantis software (mantis64.com), through a multifunction I/O acquisition board (USB-6343, National Instruments) with other input and output channels. The analog 50 Hz output trigger was saved as analog input for post hoc alignment. In the second setup (setup B), video recording, looming stimuli and optogenetic laser stimulations were triggered through Bonsai visual programming language (https://bonsai-rx.org).

### Behavioral and laser stimulation protocols

#### Experiments in arena with shelter

For experiments in the arena with shelter, mice were habituated to the new environment (7 minutes in chemogenetic experiments, 10 minutes in optogenetic experiments, and 30 minutes in fiber photometry experiments) before the first looming stimulus was presented. Looming stimulus presentation was either manually or automatically (by a custom script in Bonsai) triggered when the mouse reached a ‘threat zone’ more than 50 cm away from the shelter. Looming stimuli were presented only when the mouse faced and walked away from the shelter and was not close to a wall. The minimum inter-stimulus interval was two minutes. Sessions began with the presentation of 3 to 6 consecutive high contrast looming stimuli (99%) to sensitize the animal before decreasing the probability of presenting a looming stimulus when mice entered the threat zone to 10-25% (or 25-50% for photometry recordings of vLGN axons in SC) for the remainder of the session. Experiments with looming stimuli lasted between 30 and 180 minutes unless otherwise stated.

For optogenetic manipulation experiments in the arena with looming stimuli, laser delivery was initiated differently across experimental setups. In setup A, the laser, or a single TTL pulse in control trials, was triggered manually once the mouse had remained in the shelter for at least 5 s without showing signs of initiating shelter departure. In setup B, laser stimulation was triggered automatically based on the animal’s position within the arena. A region of interest, in this case the inside of the shelter, was manually defined to specify the spatial location at which stimulation would be triggered. Stimulation was delivered 5 s after shelter entry. During laser- on trials, the laser remained on for 30 s. Laser-on and -off conditions were alternated in a pseudo-randomized way such that each session would have a minimum of 10 Laser-on and 10 Laser-off trials. Trials were initiated in the same way for optogenetic manipulation experiments in the presence of TMT (Figure 2G-I) and experiments in the absence of threatening stimuli (Figure 2J-L). Optogenetic manipulation experiments under the effects of corticosterone were performed similarly, beginning 20 minutes after an IP injection of either corticosterone (10 mg/kg) or saline in peanut oil (Figure 3). Due to increased sensitization to looming stimuli caused by the corticosterone, intermediate-to-high contrast stimuli were used (60% and 80%, alternating). Experiments were stopped one hour after the presentation of the first stimulus.

For chemogenetic inactivation experiments with looming stimuli (Figure S3I-L), mice were placed in the escape arena 30 minutes after an IP injection of either saline or Clozapine N-Oxide (CNO, 5 mg/kg, SML2304, SIgma Aldrich). A fixed sequence of 10 looming stimuli was used (40%, 20%, 60%, 40%, 20%, 40%, 20%, 60%, 40%, 20%), presented with a probability of between 25 and 50% upon threat zone entry. The experiment was stopped after the presentation of the 10th stimulus or 50 minutes after the presentation of the first stimulus, whichever happened first.

### Elevated plus maze

Animals were allowed to explore the elevated plus maze (EPM) freely. For optogenetic manipulation experiments, the laser was manually initiated as the mouse approached the centre of a maze from a closed arm. The laser was on for 30 s during laser-on trials, while laser-off trials were initiated by manually triggering a single TTL pulse to enable behavioral alignment during analysis. Laser-on and -off conditions were alternated in a pseudo-randomized way such that each session would have, on average, a minimum of 15 laser-on and 15 laser-off trials. For chemogenetic experiments (Figure S3L), EPM sessions began 35 minutes after an IP injection of CNO (5 mg/kg). These data are reanalyzed from Fratzl et al. 2021. For corticosterone experiments (Figure 3C,H), EPM sessions began immediately after experiments with looming stimuli. All manipulation experiments were stopped 30 minutes after the first trial, regardless of trial count, to prevent habituation. Photometry experiments were stopped after 60 minutes.

### Open field and novel object exploration and other paradigms

Behavior in an open field arena was assessed 35 minutes after an IP injection of CNO (5mg/kg). Mice were allowed to freely explore the arena for ten minutes (Figure 3A,B).

For optogenetic experiments in the empty open field, mice were allowed to explore freely for one hour (Figure S5H-J). The laser or a single TTL pulse for control trials was manually triggered when the mouse performed one of four behaviors, grooming, rearing, walking, or gazing upwards. The laser stayed on for 4 s and trials alternated between laser-on trials and laser-off trials, with a minimum of 30 s between the end of one trial and the beginning of the next.

Prior to object-exploration experiments, animals were habituated to the open field arena. Twelve small objects were used, selected to be of varying colors, materials, and textures but similar sizes. In the single-object task (Figure S5A-D), animals were additionally habituated to the arena first with one object (selected at random), and then with a piece of food, placed in the middle. Throughout the test, Laser blocks, where laser stimulation automatically alternated between 30 s on and 30 s off, and Control blocks, with no laser stimulation and only a TTL pulse for alignment, were alternated for 5.5 minutes each. Object replacement took about 30 s and blocks were manually initiated for each block 30 s after object placement. In each block, animals were presented with either one of 12 novel objects, their habituated object, or a piece of food. Novel objects were always presented in the same order. Laser and Control blocks were pseudo-randomized, ensuring animals experienced both the habituated object and food twice each, in both Laser and Control conditions, separated by at least two novel objects. Single-object task experiments lasted approximately 90 minutes.

In the double-object task (Figure S5E-G), animals were presented with two novel objects at a time, in pseudo-random unique pairings: pairings were chosen prior to starting the experiment ensuring no object appeared twice in a single experiment and that all pairings are unique across experiments. Each pairing was presented for 5.5 minutes. Alternating laser or control trials, with stimulation lasting for 4 s, were triggered manually at every new object-interaction bout, with object replacement taking about 30 s and laser stimulation starting 30 s after object placement. Trials were not triggered if the interaction with an object was a continuation of, or a switch from, an interaction from the other object. Double-object task experiments lasted approximately 35 minutes.

For optogenetic manipulation experiments with the toy snake (Figures 2D-F, S4D,E) , mice were allowed to explore an arena with a toy snake for 30 minutes. Laser stimulation automatically alternated between 30 s on and 30 s off. For optogenetic experiments with a live cricket (Figure 2A-C, S4A-C), mice were allowed five minutes to explore the empty arena prior to placement of a live cricket. Each cricket and mouse pairing was unique, ensuring neither cricket nor mouse had previously experienced the other. Experiments lasted for up to 30 minutes. Laser stimulation automatically alternated between 30 s on and 60 s off.

For the place-preference test (Figure S4I,J), mice were placed in the arena for 20 minutes total: 10 minutes for habituation to the two-chamber setup and 10 minutes for optogenetic manipulation. During the 10 minutes optogenetic manipulation, laser stimulation was manually triggered upon entry to the stimulation side of the arena for the duration of the mouse’s stay on the stimulation side, or for 30 s, whichever was shorter. If the mouse stayed on the stimulation side for greater than 60 s, laser stimulation continued on a 30 s on and 30 s off protocol until the mouse returned to the non-stimulation side. Laser stimulation was always manually terminated upon the mouse’s return to the non-stimulation side.

### Visual stimuli

In Setup A visual stimuli were created and presented through custom-made scripts in MATLAB (Mathworks), using the Psychophysics Toolbox extensions^84,85^. Visual stimuli were triggered through the Mantis software (mantis64.com) and projected onto the screen. A switchable gain, Si-amplified photosensor (PDA100A2, Thorlabs) was used to record the stimuli presented on the screen for post hoc alignment. Visual looming stimuli consisted of a sequence of three dark expanding circles centered over the threat zone. The radius of the spot expanded linearly at 55 deg/s from 1 deg at onset to 20.8 deg in 360 ms. The spot was then kept at constant size for 250 ms followed by a 500 ms inter-stimulus interval. The contrast (c) of the stimulus was varied by maintaining the background luminance (I_B_) constant at 9 cd × m^−2^ and changing the luminance of the spot (I) according to a negative Weber contrast law: c=−(*I*−*IB*)/*IB*. This ensured contrasts were always defined by positive values. In Setup B, looming stimulus presentation was controlled through Bonsai, which also tracked the animal online. Each visual stimulation consisted of three consecutive expanding black spots presented over 3 s. Each spot expanded from 1° to 20° of visual angle over 330 ms and was maintained at its maximum size for 270 ms. The screen background luminance was held constant at 9 cd × m⁻².

### Optogenetic manipulation

In Setup A, laser stimulation protocols were created and presented through the Mantis software (mantis64.com). In Setup B, laser stimulation protocols were created and controlled using custom-written scripts in Bonsai and executed through a Pulse Pal pulse generator (Open Ephys). For neuronal or axonal excitation of ChR2-expressing neurons and axons, a 473 nm laser (OBIS 473 nm LX 75 mW Laser System, Coherent) was pulsed for up to 30 seconds at 20Hz using 5-ms pulses. For photoinhibition of eNpHR3.0-expressing neurons and axons, a 594 nm laser (OBIS 594 nm LS 40 mW Laser, Coherent) was continuously on for up to 30 seconds. Analog signals drove either the 473 nm laser or 594 nm laser coupled to a 200 μm fiber patch cable through an achromatic fiber port (Thorlabs). Light level was divided at the level of a rotary joint (FRJ_1x2i_FC-2FC_0.22, Doric Lenses) into two equal halves. Two identical 200 μm patch cables (MFP_200/220/900-0.37_1m_FC_SMC, Doric Lenses) were connected to the two outputs of the rotary joint and the implanted cannulas.

In Setup B, laser stimulation protocols were created and controlled using custom-written scripts in Bonsai and executed through a Pulse Pal pulse generator (Open Ephys).

Peak power was measured at the tip of the patch fibers before each experiment. The analog output driving the laser was also recorded as analog input for post hoc trial alignment. The peak laser power on each side of the laser 10 mW at the tip of each optical fiber, measured before implantation. To control for light-induced artifacts on behavior in vLGN, laser stimulation was performed in animals with GFP instead of an optogenetic construct expressed in GABAergic vLGN neurons (Figure S3E-H).

### Fiber photometry experiments

In both setups, a 473 nm LED (CLED_473, Doric Lenses) and a 405 nm LED (CLED_405, Doric Lenses) were used to excite GCaMP at its calcium-dependent and isosbestic wavelength, respectively (Lerner et al., 2015). The light was focused into a single output fiber through a six-port minicube (iFMC6_AE(405)_E1(460-490)_F1(500-540)_ E2(555-570)_F2(580-680)_S,

Doric Lenses) connecting a pigtailed rotary joint (FRJ_1x1_PT_400/430/LWMJ-0.57_1m_FCM_0.15m_FCM). A 400 μm fiber patch cable connected the rotary joint to the implanted cannula through its slim magnetic connector (MFP_400/460/1100-0.48_0.8m_FC-SMC). Peak power amplitude at the tip of the patch cord was between 500 μW and 550 μW for all experiments. An integrated fluorescence detector head (part of iFMC6, Doric) converted the detected fluorescence signals into an analog output signal.

Data was continuously acquired and synchronized through a USB-driven PyPhotometry board (Akam and Walton, 2019). The python (https://www.python.org) based open source software PyPhotometry was used to drive the 473 nm and 405 nm LEDs at roughly 100 mA each in strobing mode (Akam and Walton, 2019). The software sequentially recorded and temporally aligned the fluorescence evoked by alternating stimulation with 473 nm and 405 nm light to obtain separate time-varying traces of calcium-dependent and isosbestic GCaMP fluorescence signals. The board was also used to record the analog photometry signals at 120 Hz and the digital 50 Hz pulses driving the camera as input for post hoc alignment.

For experiments combining optogenetic manipulation with fiber photometry, PyPhotometry drove the alternating 473 nm and 405 nm LEDs and acquired the fluorescence signals at 120 Hz. Laser stimulation protocols were generated in Bonsai and executed through a Pulse Pal pulse generator (Pulse Pal v2, Sanworks). Photometry acquisition, laser stimulation and camera TTL signals were all coordinated and synchronized within Bonsai for post hoc alignment.

### Data analysis

#### Behavioral data pre-processing

In Setup A, raw videos of mouse behavior were extracted using a dedicated converter (mantis64.com) and saved as AVI files. The analog input channels (sound, photodiode, laser, and camera trigger) were extracted in MATLAB (MathWorks). All analog input channels were recorded at 25000 samples per second, while the camera was triggered at 50 Hz. Using custom-made scripts in MATLAB, time points of visual stimuli onsets, laser onsets, and TTL pulses for control trials were extracted and matched to the corresponding video camera frame.

In Setup B, raw behavioral videos and the corresponding analog and digital input channels were acquired using Bonsai. The camera was triggered at 50 Hz, and analog input channels were sampled at 50 samples per second. Using custom-written MATLAB scripts, timestamps of visual stimulus and laser onsets were extracted and aligned to the corresponding video frames.

To extract behavior we used a deep neural network approach (DeepLabCut^86^). A network was trained with 200 manually annotated frames of an initial dataset in the looming arena, and then re-trained with 300 new frames each from two additional datasets from the elevated plus maze and open field. A further 500 frames from all three paradigms were annotated and added and a final network was re-trained. The resulting labeled videos were manually inspected for quality control. This network was used for all behavioral classifications, and the XY-position of body-center, head, nose, and ears within the arenas was extracted. XY-positions were loaded and processed in MATLAB. The position of the animal was defined as the body-center position. The head-direction of the animal was computed as the vector between the nose and the head (in the arena with shelter: 0 degrees - mouse facing toward the “threat zone”; ± 180 degrees: mouse facing toward the shelter). Instantaneous speed was computed as the Euclidean distance between two positions at consecutive time points. Angular velocity was computed as the discrete derivative at two consecutive time points (taking into account circular boundary conditions). Prior to computing the speed and angular velocity, the position and head direction vectors were smoothed by a 100 ms running average filter to avoid amplifying noise when computing derivatives. The ears were tracked to increase the stability of the network with more co-dependent markers. For all experiments, data analysis was performed in custom-written routines in MATLAB (Mathworks)

### Behavior shelter leaving paradigms

Shelter exit events were defined based on a combination of criteria: animals’ body-center has crossed shelter threshold, animals’ body-center must have been inside the shelter for at least 5s beforehand, head-direction must be facing away from the shelter, and the body-center must remain outside of the shelter for at least 1 s after crossing the threshold. Since trials in optogenetics experiments were not always initiated after the same amount of time spent inside the shelter, the time to leave the shelter was computed as the time elapsed between manual trial initiation and shelter exit. Relative time spent outside of shelter was computed as time spent outside the shelter during a 60 s period following trial start in optogenetic experiments. The maximum distance from shelter was calculated as the greatest X-position of a mouse’s body-center after a shelter exit. The true possible maximum is equal to the length of the arena excluding the shelter (66 cm). In order to mimic optogenetic experimental conditions, shelter exit trials for chemogenetic inhibition experiments were defined as beginning 5 s after shelter entry. Any shelter entries where the mouse exited the shelter in under 5 s were excluded. Shelter time per entry was computed as time elapsed, in seconds, between the trial start time and the shelter exit timepoint. Relative time spent outside of shelter was computed as time spent outside the shelter as a percentage of total session time in chemogenetic experiments.

### Behavior elevated plus maze and other paradigms

For the elevated plus maze, analysis included all instances in which mice exited a close arm. An arm entry after close-arm exit was defined as the time point at which an animal entered completely (with all four limbs) into an open or closed arm from the center. Entry timepoints detected using the animals’ position as compared to the manually defined XY-coordinates of the EPM and were verified using manually annotated timepoints. The relative number of entries into an open arm was defined as the ratio between the number of entries into an open arm and the total number of exits from a closed arm. The relative time outside a closed arm was defined as the time the animal’s entire body was either in an open arm or in the exposed, central decision point, divided by the total time elapsed between each trial and the initiation of the subsequent trial.

In place-preference experiments and experiments with the toy snake, mouse location was defined as being on one side of the arena when the body-center crossed the threshold into that side and remained there for at least 1 s. Quantification of the number of approaches was performed manually. In experiments with a cricket, interactions were quantified manually. Duration of interaction time was defined as time elapsed between the timepoint at which an animal initiated contact with the cricket and termination of interaction. Behavior in the 5 s following termination of a cricket interaction is classified as an escape if the head direction changed by +-90 degrees of the head direction at the time of interaction termination with an angular velocity of > 100 degrees/s, distance travelled was at least 20 cm, and maximum speed was at least 35 cm/s. Hunting behavior was defined as sustained pursuit for at least 2 s with at least one attempted attack or bite.

In the open field test, the 40 cm by 40 cm arena was divided into a 5 by 5 grid for analysis. Using the automatically extracted positions, the amount of time spent outside of the corners, representing 84% of the open field area, was quantified and divided by the total length of recording (10 minutes). For optogenetic experiments in the open field testing action switching, behaviors (grooming, rearing, walking, and gazing upwards without rearing) were identified manually. Initiation and termination time points of these behaviors were recorded and compared to trial time points. A trial was marked as an “action-switch” trial if the mouse switched to a new behavior for at least 2 s during the trial, defined as the 5 s after trial initiation.

Behavior for all novel-object interaction experiments was annotated manually: start of object interaction was defined as the time point at which a mouse came into contact with an object. Interaction was considered terminated at the time point from which contact was stopped if the animal did not re-initiate interaction for at least 1s. In the single-object paradigm, behavior when the laser was on was compared to equivalent periods of a stimulation-free block. Interactions terminating after the end of a laser-on or control black were included in the analysis. In the double-object task, interaction outcome was annotated manually. An outcome of ‘leave’ is defined by the mouse walking away from an object it has been interacting with within 5 s of laser stimulation or control trial start and not initiating a new interaction with another object for at least 2 s afterwards. An outcome of ‘switch’ is defined by the mouse terminating interaction with one object and starting to interact with the other object within 2 s. An outcome of stay is defined by sustained interaction with the same object for at least 3 s. Relative interaction outcome is computed as the total number of each outcome as a percentage of the total annotated interactions in that session. In both paradigms, the 30 s between object placement and manual trial initiation were excluded from analysis.

### Calcium Imaging

Raw fluorescence traces were extracted and imported into MATLAB using software generously provided by Thomas Akam (https://pyphotometry.readthedocs.io/en/latest). Raw signals of calcium-dependent (GCaMP excited at with 473 nm light) and isosbestic fluorescence (GCaMP excited with 405 nm light) were filtered with a slow 10-minute running-median filter and subtracted from the original traces to correct for slow changes such as photobleaching. The isosbestic signal should not contain any calcium-dependent signals, but is sensitive to transient changes in fluorescence not due to changes in the calcium concentration, for instance signals due to motion artifacts such as bending of or tension within the patch-cord. Such signals are shared between the two channels. To correct for such non-calcium dependent signals and artifacts, the isosbestic fluorescence trace was linearly fitted to the calcium-dependent GCaMP fluorescent signal and subtracted, providing a measure of relative change in fluorescence (ΔF/F). To be able to compare calcium activity across sessions and mice, z-scored ΔF/F was computed by subtracting the mean value of the motion-corrected calcium-dependent GCaMP signal of the session and dividing the resulting trace by the standard deviation. Fluorescence traces, acquired at 120 Hz, were then re-sampled at 50 Hz, and aligned to camera frames via the digital input signal. Displayed fluorescent traces are down-sampled to 25 Hz.

To assess differences in vLGN activity depending on recency of visual threat experience, shelter leaving events immediately following a looming stimulus-triggered escape were compared to the all shelter leaving events after the second post-escape shelter exit (Figure S1B). To determine if ramping vLGN activity before shelter exit was caused by locomotion in the shelter before exit, we divided trials into two groups: trials where movement began over 4s prior to shelter exit, and trials where movement began under 2 s prior to shelter exit (Figure S1C). Movement bouts inside the shelter (Fig S1D) were defined as sudden initiation of sustained movement while in the shelter for greater than 5 s not resulting in a shelter exit following a period of at least 2 s spent stationary.

### Decoding of neural activity before leaving safety

To quantify whether exits from safety could be predicted from vLGN calcium activity, we trained a supervised classifier using windows of peri-event calcium activity preceding shelter exits in the shelter leaving paradigm or events of leaving a closed arm on the elevated plus maze. For each leaving event, a fixed temporal window spanning from 3.0 s before until time of shelter or closed arm exit was extracted from the normalized photometry signal (sampling rate 50 Hz; 150 samples per window). Each time bin within the window was treated as an independent feature, yielding a multivariate representation of activity dynamics leading up to the event of leaving safety. A binary random forest classifier (TreeBagger; 300 trees) was trained to discriminate these activity periods preceding exits from safety from random control periods. Control periods of the same length were randomly selected from the same recording session, matched in number to the leaving events, and at least 2.0 s away from any leaving safety event to avoid temporal overlap.

For each animal, class-balanced windows were split 70%/30% (stratified) into training and held-out test sets. From held-out predictions we report accuracy (% correctly classified at P(LeaveSafety) = 0.5 threshold) and row-normalized confusion matrices (per animal, averaged across animals), which express the conditional probability of predicted class given true class and are not the same computation as accuracy, though related for balanced test sets.

Significance was assessed against a shuffled null: for each comparison, class labels were randomly permuted and a classifier retrained on the identical train/test split, repeated 200 times to build a null accuracy distribution. Group significance used a one-tailed Wilcoxon signed-rank test (real vs. shuffled accuracy across animals) and a Stouffer-combined p-value across animals’ individual permutation tests.

To test cross-paradigm generalization, the shelter-leaving-trained classifier was applied without retraining to EPM closed-arm-exit and control windows (same window/feature structure). Significance used the same permutation approach: each null model was retrained on shuffled shelter-leaving labels and applied, without retraining, to the EPM data, giving a transfer null distribution comparable to the real transfer accuracy.

In addition to within-animal decoding, a pooled decoder was trained and evaluated for each paradigm using trials pooled across all animals, with a single random, class-stratified 70%/30% trial-level split defining training and test sets (Figure S2B,D). Unlike the leave-one-animal-out analysis, this pooled decoder’s training and test sets can both include trials from the same animal, evaluating whether a shared decodable signal exists across the pooled population rather than generalization to a specific unseen animal. For both within-animal and pooled decoders, we additionally examined the distribution of held-out predicted probabilities, P(LeaveSafety), separately for true leaving-safety and true control trials, to visualize class separation underlying the reported accuracy and confusion matrices.

### Statistics

All statistical analyses were performed using MATLAB. Unless otherwise stated, data are presented as mean ± SEM across mice, with faint dots showing individual animals. Fiber photometry traces show the mean across mice, with shaded areas indicating SEM. Trial-level measures (for example, time to leave the shelter, or the probability of an open-arm entry) were first averaged within each mouse.

For comparisons between two conditions, paired tests were used for within-animal comparisons. These included laser versus no laser (control) trials within the same session and comparisons between conditions in the same animal. We used paired t-tests, or Wilcoxon signed-rank tests where data were not normally distributed, as assessed by the Shapiro–Wilk test. Comparisons between two independent groups of mice used two-tailed unpaired Student’s t-tests for normally distributed data, Welch’s t-test where variances were unequal, or Mann– Whitney U tests for non-normally distributed data. All tests were two-tailed except where stated in the figure legends. One-tailed tests were used for the experiments in Figures 2 and S4, based on the a priori directional hypothesis from Figure 1 that vLGN activation would increase approach and exploration. Early versus late session epochs were tested separately.

Where three conditions were compared, an omnibus test was performed before pairwise comparisons. When all three groups consisted of independent cohorts of mice (Figures 3B,C and S6B,C), we used a one-way ANOVA followed by Tukey’s post hoc test. Normality of residuals was assessed with the Shapiro-Wilk test and homogeneity of variance with Levene’s test. When the design combined an independent control cohort with two conditions tested within the same animals (saline mice vs. +Cort mice tested with and without laser; Figures 3G,H and S6D,E,G), we used a linear mixed-effects model with condition as a fixed effect and animal as a random intercept. The effect of condition was assessed with an F-test using Kenward–Roger degrees of freedom, followed by pairwise comparisons with Holm correction for multiple comparisons. Where one group showed no variance (Figure S6F), pairwise comparisons were instead made with Mann–Whitney U tests (independent group) or Wilcoxon signed-rank tests (within-animal), with Holm correction.

No statistical methods were used to predetermine sample sizes. Sample sizes were similar to those used in previous studies with comparable paradigms^30,43^. Animals were excluded from analysis if post hoc histology showed off-target viral expression or fiber placement outside the target region.

