## Supplementary Figures for "Prethalamus integrates neocortical and hypothalamic input to promote courage"

### Supplemental figures

#### Figure S1, related to Figure 1

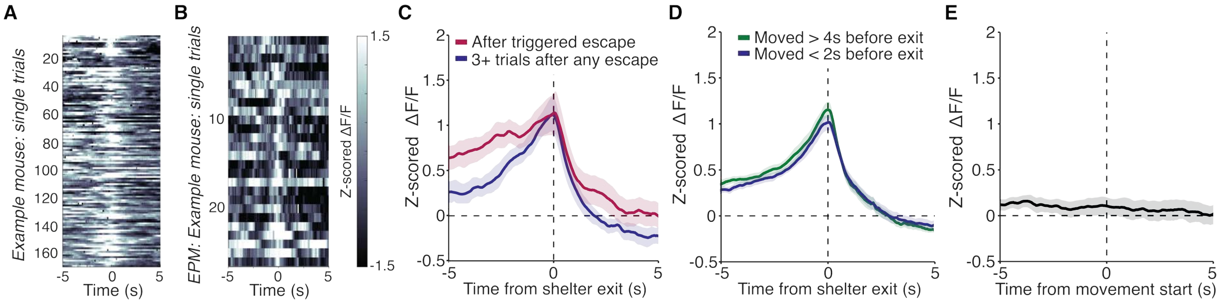


**Figure S1. Calcium activity of GABAergic vLGN axons in mSC around leaving safety in different conditions**.

(A) Calcium activity of GABAergic vLGN axons in mSC of an example mouse during individual leaving shelter events (trials) aligned to shelter exit. (B) Calcium activity of GABAergic vLGN axons in mSC of an example mouse on the elevated plus maze (EPM) during individual events of leaving a close arm, aligned to closed arm exit. (C) Mean calcium activity of GABAergic vLGN axons in mSC aligned to shelter exit for shelter leaving events immediately following a looming stimulus-triggered escape (red) or leaving shelter events after the animals has left the shelter at least two times before without escapes (blue). N = 15 mice, pale shading represents SEM across mice. (D) Mean calcium activity of GABAergic vLGN axons in mSC aligned to shelter exit in trials in which animals started to locomote more than 4 s before shelter exit (green), and when animals initiated locomotion less than 2 s prior to shelter exit (blue). N = 15 mice. (E) Mean calcium activity of GABAergic vLGN axons in mSC aligned to animals' movement onset in the shelter for movement bouts in which the mouse did not exit the shelter for at least 5s. N = 15 mice, pale shading represents SEM across mice.

#### Figure S2, related to Figure 1

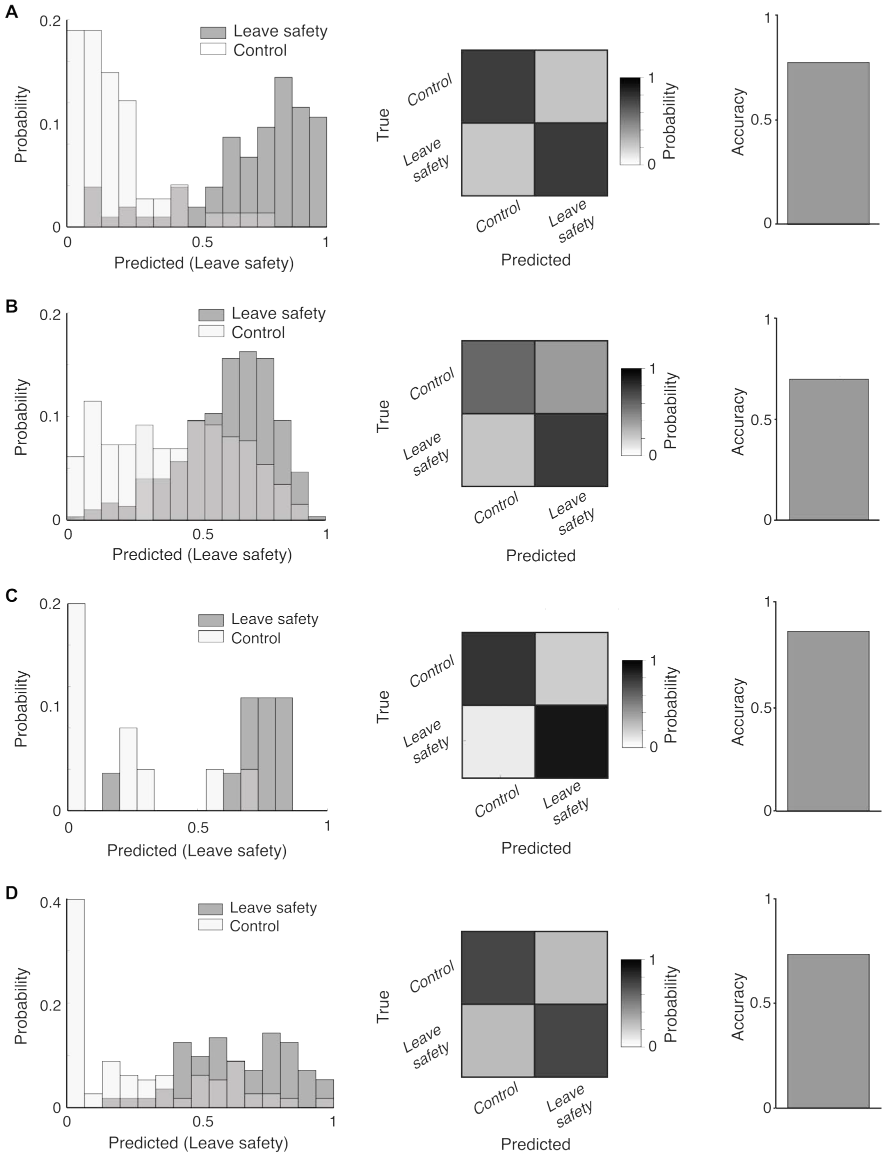


**Figure S2. Example decoding performance and pooled decoder across animals**.

(A) Decoder performance for a representative animal in the shelter leaving paradigm. Left, distribution of predicted probabilities that a period of calcium activity preceded shelter exit or not for calcium activity -3 to 0s before shelter exit (dark grey bars) and random 3s control periods (light grey bars) in held out data. Middle, row-normalized confusion matrix showing classification performance between control periods and shelter exit events. Right, classification accuracy for this animal. (B) Same as A, but for pooled decoding performance for the leaving shelter paradigm; training and testing was performed on trials pooled across all animals. (C) Same as A, but for calcium activity from a representative animal on the elevated plus maze, decoding closed arm exits versus random control periods. (D) Same as B, but for pooled decoding performance across animals on the elevated plus maze, decoding closed arm exits versus random control periods.

#### Figure S3, related to Figure 1

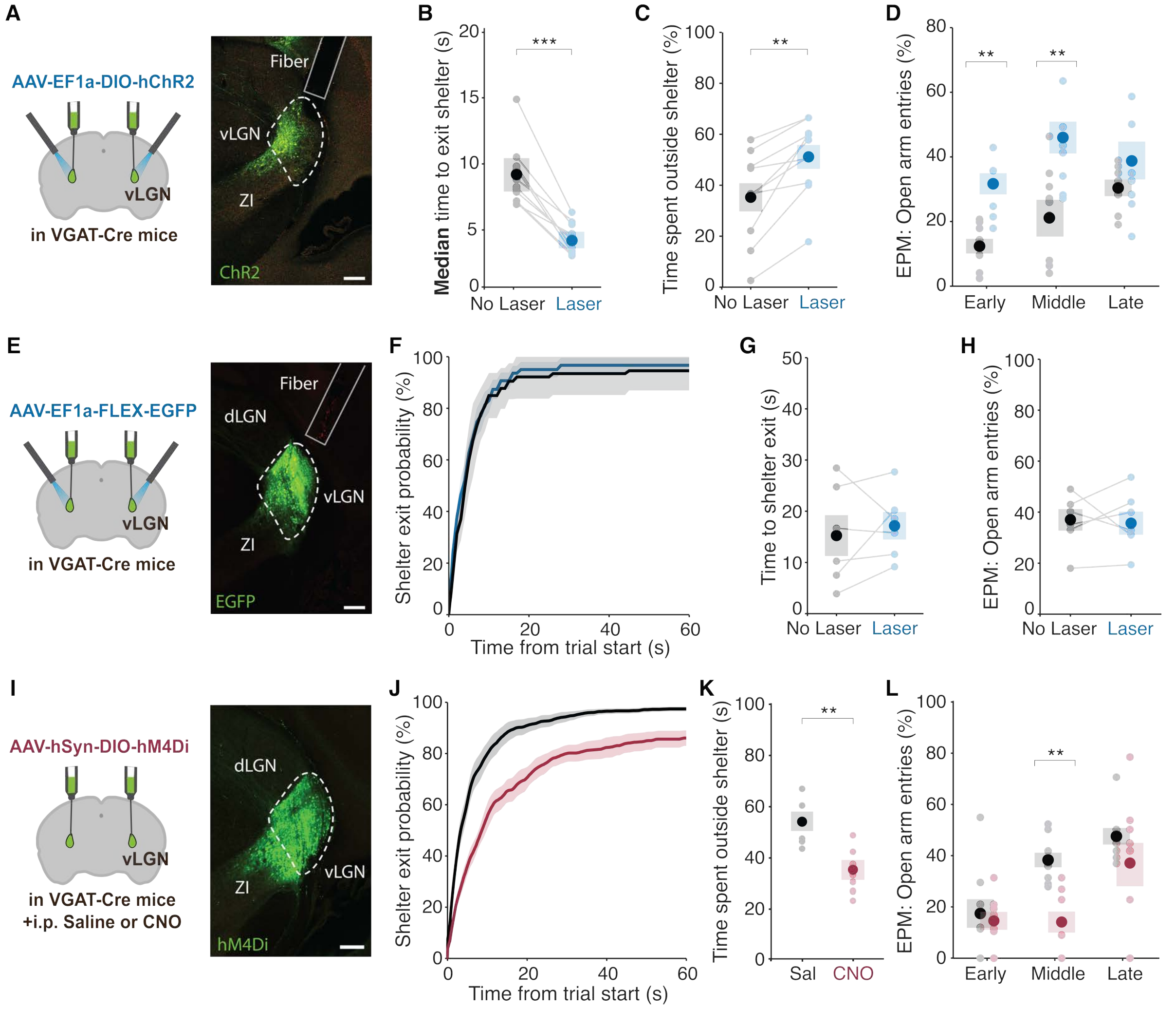


**Figure S3, Optogenetic activation and chemogenetic inactivation of GABAergic vLGN neurons and controls.**

(A) Left, experimental approach. Right, EYFP-expressing GABAergic vLGN neurons and outlined fiber track. Scale bar = 200 µm. (B) Median time to exit shelter during vLGN activation (blue) and in control trials (black). P < 0.001, two-tailed paired t-test, N = 10 mice. Same data as in Figure 1J, but computing median instead of mean time to shelter exit for each animal. (C) Time spent outside of the shelter as a percentage of total trial time (60s) during optogenetic activation of GABAergic vLGN neurons (blue) and in control trials (black). P = 0.0027, two-tailed paired t-test, N = 10 mice. (D) Probability of an EPM open arm entry from a closed arm during vLGN activation (blue) and in control trials (black) in the first 10 minutes (early, P = 0.0038, two-tailed paired t-test), between 11 and 20 minutes (middle, P = 0.0066, two-tailed paired t-test) and in the last 10 minutes on the EPM (late, P = 0.503, two-tailed paired t-test), N = 10 mice. (E) Left, experimental approach. Right, GFP-expressing GABAergic vLGN neurons and outlined fiber track. Scale bar = 200 µm. (F) Mean cumulative probability of animals' exit from shelter over 60s in trials with laser stimulation of GFP-expressing GABAergic vLGN neurons (blue) and in control trials without laser (black) within the same session. N = 6 mice. (G) Mean time until mice leave the shelter after trial start during laser stimulation (blue) and in control trials (black). P = 0.581, two-tailed paired t-test, N = 6 mice. (H) Probability of an EPM open arm entry in laser stimulation (blue) and control trials (black). P = 0.610, two-tailed paired t-test, N = 6 mice. (I) Left, experimental approach. Right, hM4Di-expressing GABAergic vLGN neurons. Scale bar = 200 µm. (J) Mean cumulative probability of animals' exit from the shelter over 60s in mice with chemogenetic inactivation of GABAergic vLGN neurons (red, N = 7 mice) and control mice (black, N = 6 mice). (K) Mean time spent outside the shelter in mice with chemogenetic vLGN inactivation (red, N = 7 mice) and control mice (black, N = 6 mice). P = 0.0048, unpaired Student's t-test. (L) Probability of an EPM open armentry in mice approaching the center from a closed arm with chemogenetic vLGN inactivation (red) and in control mice (black) in the first 10 minutes (early, P = 0.444, unpaired Student's t-test), between 11 and 20 minutes (middle, P = 0.0029, unpaired Student's t-test), and in the last 10 minutes on the EPM (late, P = 0.943, Mann-Whitney U test). In all plots, faint dots represent data from individual mice. Dark dots and error bars represent mean and SEM across mice.

#### Figure S4, related to Figure 2

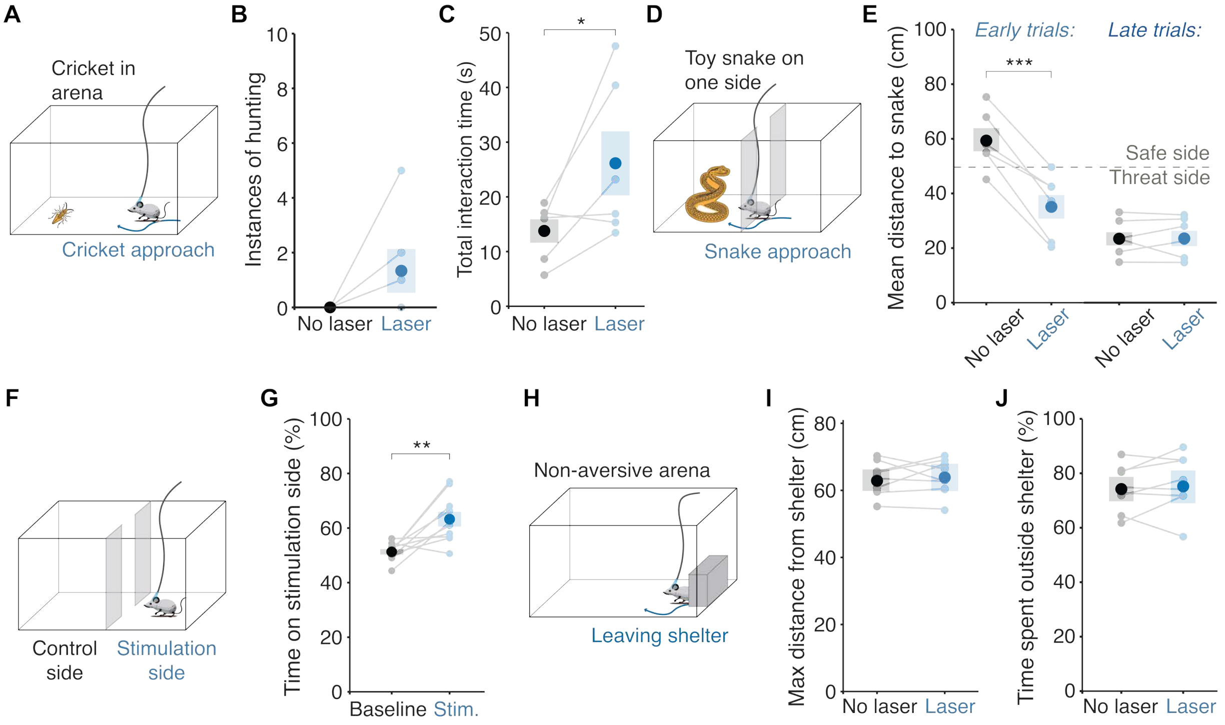


**Figure S4, vLGN activation promotes threat-discounting behavior.**

(A) Schematic of the experimental arena with a live cricket. (B) Number of instances of hunting during activation of ChR2-expressing GABAergic vLGN neurons (blue) and in control periods (black). P = 0.25, Wilcoxon signed-rank test, N = 6 mice. (C) Total time spent interacting with the cricket during vLGN activation (blue) and in control periods (black). P = 0.0275, one-tailed paired t-test. (D) Schematic of experimental arena with dark walls separating two halves of the arena and a toy snake on one side. (E) Mean distance between mice and the snake during optogenetic activation of ChR2-expressing GABAergic vLGN neurons (blue) and in control periods (black), early (left, first 5 minutes; P = 0.0007, one-tailed paired t-test, N = 6 mice) and late in the experiment (right, last 5 minutes; P = 0.913). (F) Schematic of non-aversive arena with shelter. (G) Maximum distance reached from shelter with no threat stimuli after shelter exit with (blue) and without vLGN activation (black). P = 0.642, one-tailed paired t-test, N = 8 mice. (H) Percentage of time spent outside of the shelter during optogenetic activation of GABAergic vLGN neurons (blue) and during control trials (black). P = 0.196, one-tailed paired t-test, N = 8 mice. (I) Schematic of place-preference test. (J) Time spent on stimulation side in the baseline condition (10-minute period without laser, black) and when GABAergic vLGN neurons were activated whenever mice entered the stimulation side (10-minute period with laser stimulation on one side, blue). P = 0.0028, one-tailed paired t-test, N = 10 mice. In all plots, faint dots represent data from individual mice. Dark dots and error bars represent mean and SEM across mice.

#### Figure S5, related to Figure 2

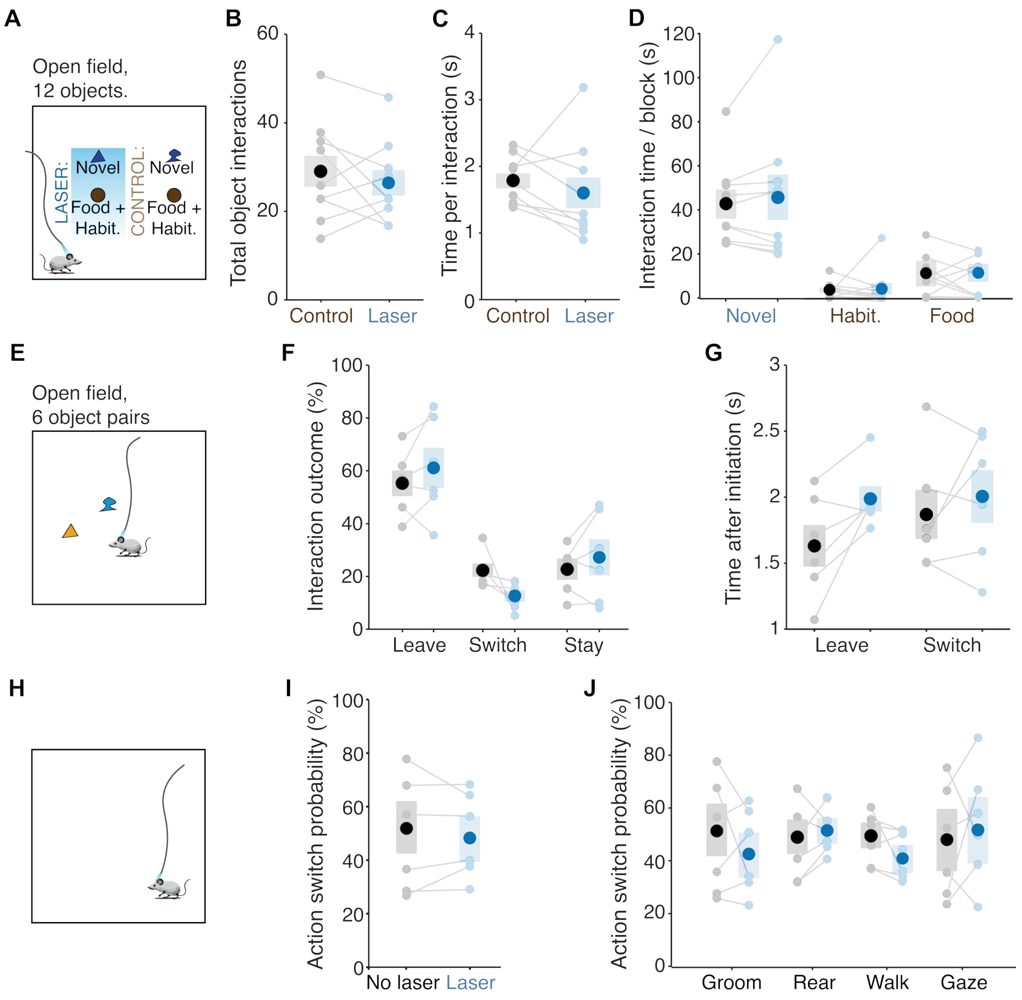


**Figure S5, vLGN activation does not affect behavior outside of anxiogenic contexts.**
(A) Schematic of experimental protocol: a sequence of individual objects (one of 10 novel objects, food or a habituated object) are placed in an open field during alternating 5-minute laser stimulation or control periods. (B) Total number of novel object interactions, during 5-minute optogenetic activation period of GABAergic vLGN neurons (blue) or in control blocks without laser stimulation (black). P = 0.392, paired two-tailed t-test. N = 9 mice. (C) Mean duration of object interactions during vLGN activation (blue) or in control blocks (black). P = 0.191, Wilcoxon signed-rank test. N = 9 mice. (D) Mean total time spent interacting with novel objects, habituated objects, and food per 5-minute period during vLGN activation (blue) or in control blocks (black). All P-values > 0.05, Wilcoxon signed-rank test. N = 9 mice. (E) Schematic of experimental protocol: 2 novel objects were presented simultaneously, and laser stimulation or control trials were triggered at the beginning of new interaction bouts. (F) Outcome of novel object interaction within 5s of starting vLGN activation (blue) or control trials (black), remain with the object (P = 0.288, paired two-tailed t-test), switch to the other object (P = 0.0741) or leave the object without switching (P = 0.288), N = 6 mice. (G) Average time until mice left the object (P = 0.05, paired two-tailed t-test) or switched to the other object (P = 0.448) after initiation of vLGN activation (blue) or control trials (black), N = 6 mice. (H) Experimental protocol: free exploration in an empty arena. The laser was triggered during different actions. (I, J) Probability of mice changing their current action within 5s of initiation of vLGN activation (blue) or control trials (black) for all actions combined (I, P = 0.411, paired two-tailed t-test) and for specific actions (J) including grooming (P = 0.762, paired two-tailed t-test), rearing (P = 0.839), walking (P = 0.094), and gazing upwards (P = 0.429), N = 6 mice. In all plots, faint dots represent data from individual mice. Dark dots and error bars represent mean and SEM across mice.

### Figure S6, related to Figure 3

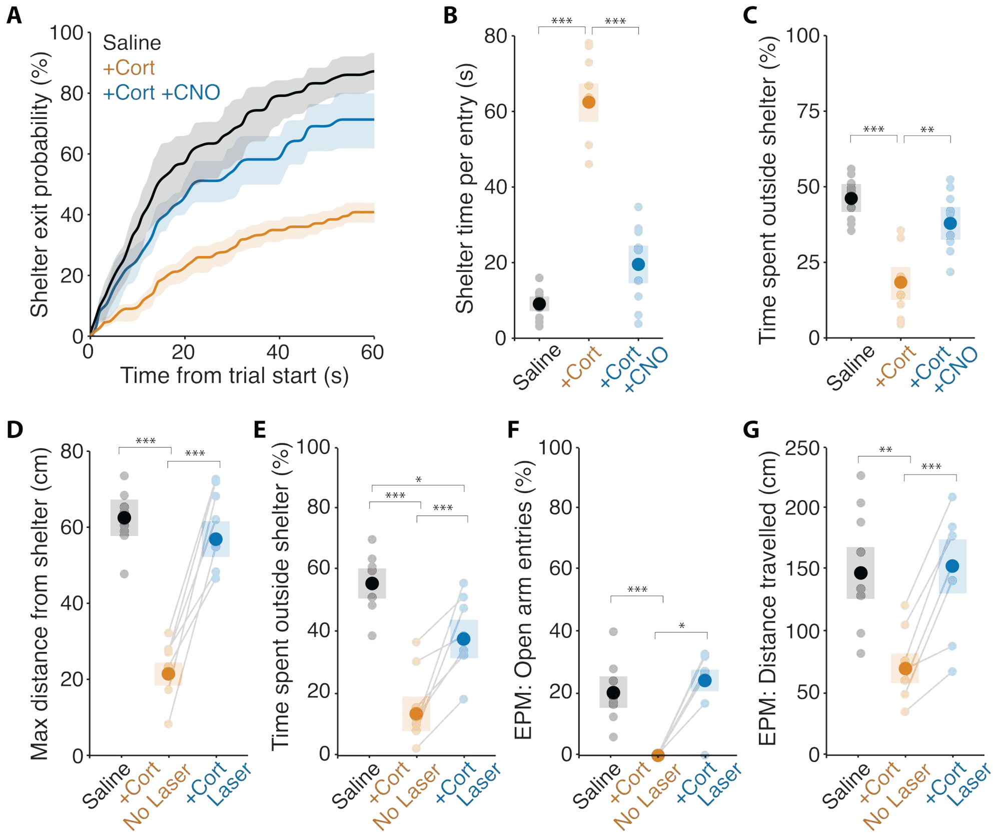


**Figure S6. vLGN activation rescues stress-induced anxiety-like behavior.**
(A) Cumulative probability of animals' exit from shelter within 60 s of trial start, in mice expressing excitatory DREADDs (hM3D), injected with saline (black, N = 8 mice), with corticosterone (+Cort, brown, N = 8 mice), or with corticosterone and CNO (+Cort+CNO, blue, N = 8 mice). (B) Time spent inside the shelter per entry. One-way ANOVA, $F(2,21) = 80.2$, $P < 0.0001$, followed by Tukey's post hoc test: saline vs +Cort, $P < 0.0001$; saline vs +Cort+CNO, $P = 0.12$; +Cort vs +Cort+CNO, $P < 0.0001$. (C) Time spent outside the shelter as a percentage of total session time. One-way ANOVA, $F(2,21) = 16.24$, $P < 0.0001$, followed by Tukey's post hoc test: saline vs +Cort, $P < 0.0001$; saline vs +Cort+CNO, $P = 0.23$; +Cort vs +Cort+CNO, $P = 0.0025$. (D) Maximum distance away from shelter during the 60 s after trial start in saline mice (black, N = 8 mice) and in corticosterone-injected mice (+Cort, N = 8 mice) without (brown) and with (blue) optogenetic activation of GABAergic neurons in vLGN. Linear mixed-effects model with condition as fixed effect and animal as random intercept (Kenward-Roger degrees of freedom), $F(2,11.8) = 248.9$, $P < 0.0001$, followed by Holm-corrected pairwise comparisons: saline vs +Cort No Laser, $P < 0.0001$; saline vs +Cort Laser, $P = 0.50$; +Cort No Laser vs +Cort Laser, $P < 0.0001$. (E) Percentage of time spent outside the shelter in the same conditions. Statistics as in D: $F(2,11.8) = 84.0$, $P < 0.0001$; saline vs +Cort No Laser, $P < 0.0001$; saline vs +Cort Laser, $P = 0.012$; +Cort No Laser vs +Cort Laser, $P < 0.0001$. (F) Percentage of times mice chose to enter the open arm after approaching the central choice point from a closed arm on the EPM in saline mice (black, N = 8 mice) and in +Cort mice (N = 8 mice) without (brown) and with (blue) optogenetic activation of GABAergic neurons in vLGN at the central choice point. Because no +Cort No Laser mouse entered the open arm, comparisons were made with Mann–Whitney U tests (saline vs +Cort conditions) and a Wilcoxon signed-rank test (+Cort No Laser vs +Cort Laser), with Holm correction: saline vs +Cort No Laser, $P = 0.0005$; saline vs +Cort Laser, $P = 0.40$; +Cort No Laser vs +Cort Laser, $P = 0.031$. (G) Distance travelled on the EPM in the same conditions. Statistics as in D: $F(2,11.8) = 33.4$, $P < 0.0001$; saline vs +Cort No Laser, $P = 0.005$; saline vs +Cort Laser, $P = 0.85$; +Cort No Laser vs +Cort Laser, $P = 0.0002$. In all plots, faint dots represent data from individual mice. Dark dots and error bars represent mean and SEM across mice.

#### Figure S7, related to Figure 4

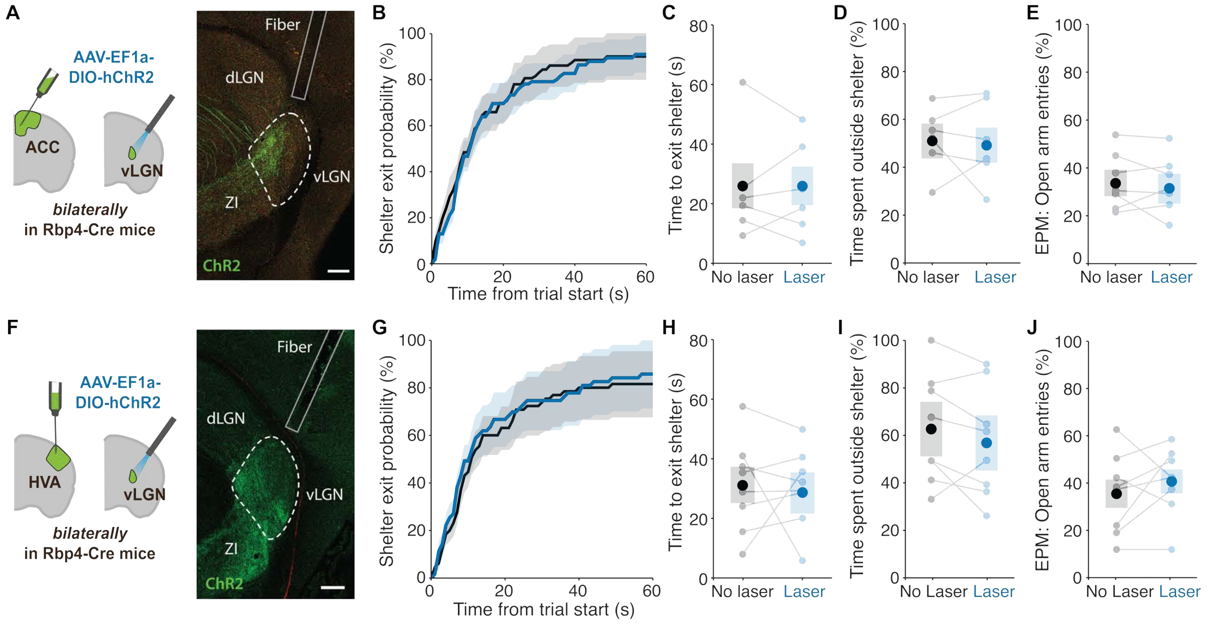


**Figure S7, Anterior cingulate cortex and higher visual area inputs to vLGN do not affect decisions to leave safety.**

(A) Left, schematic of bilateral experimental approach. Right, representative image of ChR2-expressing ACC axons in vLGN and outlined fiber track. Scale bar = 200 µm. (B) Cumulative probability of animals' exit from shelter within 60s of trial start during optogenetic activation of anterior cingulate cortex (ACC) axons in vLGN (blue) and control trials (black) within the same session. N = 6 mice. (C) Mean time until mice leave shelter after trial start during ACC axon activation (blue) and control trials (black). P = 0.877, paired two-tailed t-test, N = 6 mice. (D) Percentage of time spent outside of shelter during ACC axon activation (blue) and control trials (black). P = 0.717, paired two-tailed t-test, N = 6 mice. (E) Probability of entering an open arm on the EPM from the central choice point coming from a closed arm as a percentage of total number of trials with optogenetic activation of ACC axons (blue) and of control trials (black). P = 0.781, paired two-tailed t-test, N = 6 mice. (F) Left, schematic of bilateral experimental approach. Right, representative image of ChR2-expressing lateral higher visual area (HVA) axons in vLGN and outlined fiber track. Scale bar = 200 µm. (G) Cumulative probability of animals' exit from shelter within 60s of trial start during optogenetic activation of HVA axons in vLGN (blue) and control trials (black) within the same session. N = 8 mice. (H) Mean time until mice leave shelter after trial start during HVA axon activation (blue) and control trials (black). P = 0.939, N = 8 mice. (I) Percentage of time spent outside of the shelter during HVA axon activation (blue) and control trials (black). P = 0.280, paired two-tailed t-test, N = 8 mice. (J) Probability of entering an open arm on the EPM from the central choice point coming from a closed arm as a percentage of total number of trials with optogenetic activation of HVA axons (blue) and control trials (black). P = 0.407, paired two-tailed t-test, N = 8 mice. In all plots, faint dots represent data from individual mice. Dark dots and error bars represent mean and SEM across mice.

#### Figure S8, related to Figure 4

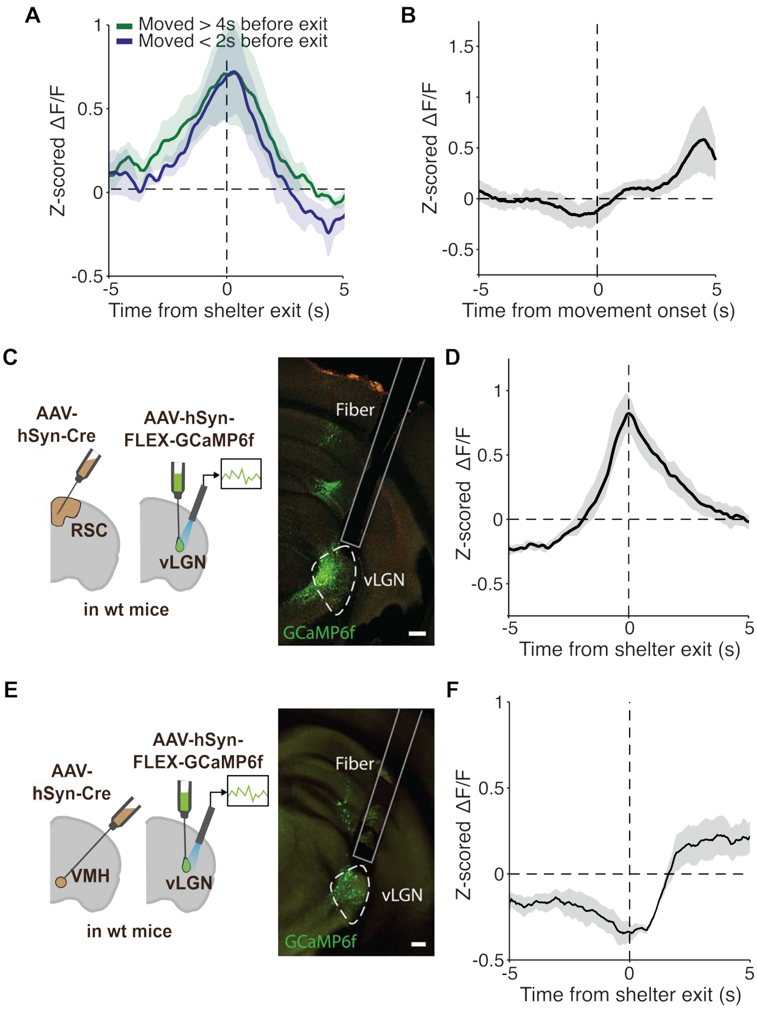


**Figure S8**, **Additional characterization of RSC inputs to vLGN and responses of vLGN neurons receiving different inputs.**

(A) Mean calcium activity of retrosplenial cortex (RSC) axons in vLGN aligned to animals leaving shelter when animals were locomoting within the shelter for more than 4 seconds before exit (green) and when animals initiated locomotion less than 2 seconds prior to shelter exit (blue). N = 8 mice, pale shading represents SEM across mice. (B) Mean calcium activity of RSC axons in vLGN aligned to animals' movement onset in shelter for movement bouts in which the mouse did not exit the shelter for at least 5s. N = 8 mice, pale shading represents SEM across mice. (C) Left, schematic of experimental approach. Right, representative image shows labelled RSC-innervated vLGN somas and fiber placement. Scale bar = 200 µm. (D) Mean calcium activity from fiber photometry of RSC-innervated vLGN neurons aligned to shelter exit. N = 8 mice, pale shading represents SEM across mice. (E) Left, schematic of experimental approach. Right, representative image shows labelled ventromedial hypothalamus (VMH)-innervated vLGN somas and fiber placement. Scale bar = 200 µm. (F) Mean calcium activity from fiber photometry of VMH-innervated vLGN neurons aligned to shelter exit. N = 6 mice, pale shading represents SEM across mice.

#### Figure S9, related to Figure 4

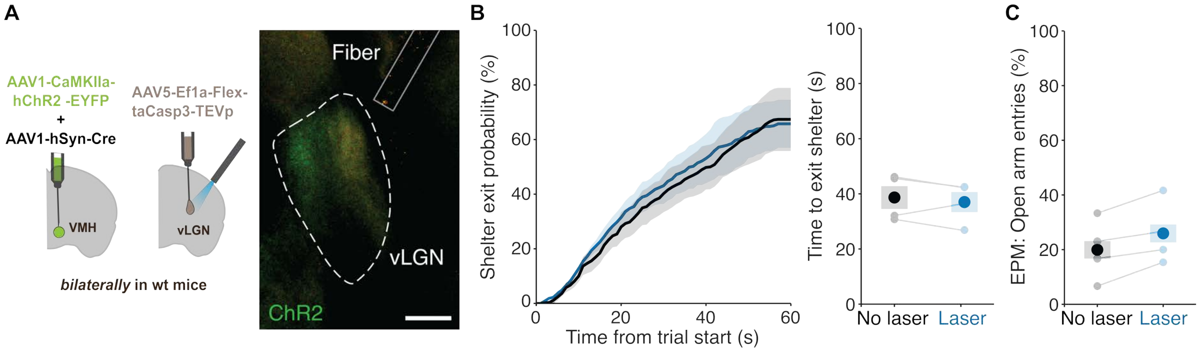


**Figure S9, Projection-specific ablation VMH-innervated vLGN neurons prevents laser stimulation effect.**

(A) Left, schematic of bilateral experimental approach with Cre dependent taCasp3 expression in the vLGN. Right, representative image shows labelled VMH axons and fiber placement in the vLGN. Scale bar = 200 µm. (B) Left, cumulative probability of animals' exit from shelter within 60s of trial start during optogenetic activation of VMH axons in vLGN (blue) and control trials (black) within the same session. N = 4 mice. Right, mean time until mice leave shelter after trial start. P = 0.7769, two-tailed paired t-test. N = 4 mice. (C) Probability of entering an open arm on the EPM from the central choice point coming from a closed arm as percentage of total number of trials with optogenetic activation of VMH axons (blue) and control trials (black). P = 0.4828, two-tailed paired t-test. N = 4 mice. In all plots, faint dots represent data from individual mice. Dark dots and error bars represent mean and SEM across mice.

#### Figure S10, related to Figure 5

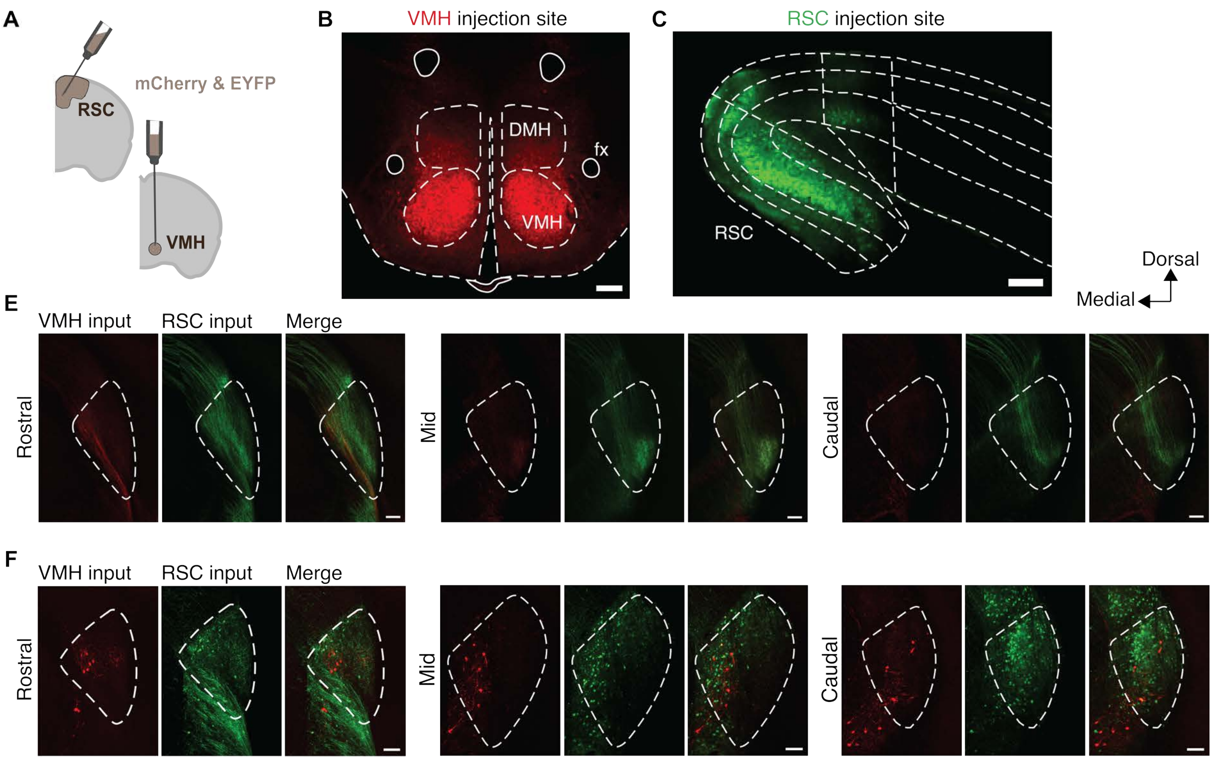


**Figure S10**, **Anatomical organization of VMH and RSC inputs in the vLGN.**

**(**A) Schematic of the dual injection approach, with mCherry and EYFP expressing viruses injected into the RSC and VMH. (B) Representative image of the VMH injection site (red). DMH, dorsomedial hypothalamus; fx, fornix. Scale bar = 100 µm. (C) Representative image of the RSC injection site (green). Scale bar = 100 µm. (D) Representative images of VMH (red) and RSC (green) axons in the vLGN (dashed outline) at rostral, mid and caudal levels, shown separately and merged. Scale bars = 100 µm. (E) Representative images of vLGN cells receiving input from VMH (green) and RSC (red) labelled with the approach shown in Figure 5B, at rostral, mid and caudal levels, shown separately and merged. Scale bars = 100 µm.
